# Polycomb Repressive-Deubiquitinase Complex Safeguards Oocyte Epigenome and Female Fertility by Restraining Polycomb Activity

**DOI:** 10.1101/2025.07.24.666633

**Authors:** Jinwen Kang, Peiyao Liu, Shoko Ichimura, Lauryn Cook, Zhiyuan Chen

**Affiliations:** Reproductive Sciences Center, Division of Developmental Biology, Cincinnati Children’s Hospital Medical Center, Cincinnati, OH 45229, USA; Department of Pediatrics, University of Cincinnati College of Medicine, Cincinnati, OH 45267, USA

**Keywords:** PR-DUB, BAP1, H3K27me3, H2AK119ub1, H3K27ac, oocyte, female fertility

## Abstract

Mouse oocytes exhibit a unique chromatin landscape characterized by broad H3K27ac and H3K27me3 domains, demarcating euchromatin and facultative heterochromatin, respectively. However, the mechanisms underlying this non-canonical landscape remain elusive. Here we report BAP1, a core component of the Polycomb Repressive-Deubiquitinase (PR-DUB) complex, as a key negative regulator of Polycomb activity during oogenesis. BAP1 restricts pervasive H2AK119ub1 accumulation in oocytes and protects oocyte-specific broad H3K27ac, particularly within gene-poor regions, from ectopic H3K27me3 deposition. While PR-DUB has been linked to gene repression, in oocytes BAP1 primarily promotes transcription and contributes minimally to Polycomb-mediated silencing. BAP1-dependent transcriptional activation is essential for oocyte developmental competence and female fertility. BAP1 loss disrupts the maternal-to-zygotic transition and impairs embryonic enhancer activation, ultimately compromising preimplantation development. Notably, while H3K27ac patterns are reset after fertilization, the aberrant H3K27me3 landscape established in BAP1-deficient oocytes persists in early embryos. Together, these findings reveal a critical role for PR-DUB in safeguarding the oocyte epigenome by protecting euchromatin from ectopic Polycomb activity, rather than enforcing transcriptional repression.

## Introduction

Polycomb group (PcG) proteins play a central role in establishing and maintaining epigenetic silencing during animal development (Kim and Kingston, 2022; Piunti and Shilatifard, 2021). PcG proteins assemble into two major complexes, Polycomb Repressive Complex 1 (PRC1) and PRC2, which function cooperatively to mediate transcriptional repression (Blackledge and Klose, 2021; Tamburri et al., 2024). In mammals, PRC1 contains the E3 ubiquitin ligases RING1A and RING1B, which catalyze monoubiquitination of histone H2A at lysine 119 (H2AK119ub1) (de Napoles et al., 2004; Wang et al., 2004). The PRC2 comprises the methyltransferases EZH1 and EZH2, which catalyze mono-, di-, and tri-methylation of histone H3 at lysine 27 (H3K27me1/2/3) (Cao et al., 2002; Kuzmichev et al., 2002; Muller et al., 2002). These complexes are typically recruited to promoters of key developmental regulators where they deposit H2AK119ub1 and H3K27me3, hallmarks of facultative heterochromatin (Margueron and Reinberg, 2011; Schuettengruber et al., 2017). Collectively, PRC1 and PRC2 mediate epigenetic silencing by promoting chromatin compaction (Francis et al., 2004) and counteracting transcription (Szczurek et al., 2024).

Oogenesis provides a unique *in vivo* model to investigate chromatin-mediated gene regulation, as postnatal oocytes grow for extended periods of time without undergoing cell division. Shortly after birth, oocytes are enclosed in primordial follicles and arrested at the diplotene stage of meiotic prophase I, known as the germinal vesicle (GV) stage. After primordial follicle activation, they enter growth phase to progress from primary follicles to pre-antral follicles, and pre-ovulatory follicles. During this growth phase (∼3-weeks in mouse), GV oocytes undergo significant cytoplasmic and nuclear enlargement without cell cycle progression and ultimately mature into fully grown GV oocytes (FGOs). During this time, oocytes accumulate maternal RNAs and proteins essential for meiotic resumption and early embryonic development (Schultz et al., 2018).

In parallel with transcriptional and translational changes, oocytes acquire a distinct epigenome, including the formation of broad, non-canonical domains of H2AK119ub1 and H3K27me3 (Chen et al., 2021; Liu et al., 2016; Mei et al., 2021; Zheng et al., 2016; Zhu et al., 2021). Notably, PRC1 and PRC2 have different functions during postnatal oocyte development. Loss of PRC1 in oocytes leads to widespread gene de-repression and disrupts the organization of Polycomb-associated three-dimensional chromatin domains (Du et al., 2020; Posfai et al., 2012). As a result, PRC1-null oocytes fail to support embryonic development beyond the two-cell stage (Posfai *et al*., 2012). In contrast, maternal loss of PRC2 has only a modest effect on oogenesis and primarily impacts non-canonical genomic imprinting dependent on maternal H3K27me3, with developmental consequences emerging after implantation (Chen et al., 2019; Chen and Zhang, 2020; Inoue, 2023; Inoue et al., 2018; Inoue et al., 2017; Matoba et al., 2022; Oberin et al., 2024). Despite these insights, how broad Polycomb domains are established and maintained during oocyte growth remains unclear. In particular, the mechanisms that restrict the spread of facultative heterochromatin into euchromatic regions during this period are poorly understood.

In addition to Polycomb domains, FGOs exhibit pervasive acetylation of histone H3 at lysine 27 (H3K27ac), a marker of active euchromatin (Liu et al., 2024; Wang et al., 2022). This non-canonical, widespread H3K27ac is mutually exclusive with H3K27me3 domains, spatially segregating euchromatin and facultative heterochromatin in oocytes (Burton and Torres-Padilla, 2025). The broad H3K27ac feature is conserved in human oocytes (Wu et al., 2023). Remarkably, many FGO-specific putative enhancers, marked by distal H3K27ac, reside in gene-poor regions, and are linked to oocyte-specific gene activation (Liu *et al*., 2024). However, the mechanisms governing the formation and function of these oocyte-specific broad H3K27ac domains remain largely unknown. These observations highlight the current knowledge gaps of how chromatin regulators coordinate the formation of spatially distinct active and repressive domains during oocyte growth.

The ubiquitin carboxy-terminal hydrolase BAP1 is the catalytic subunit of the Polycomb Repressive-Deubiquitinase (PR-DUB) complex, which removes monoubiquitin from H2AK119ub1 (Ge et al., 2023; Scheuermann et al., 2010; Thomas et al., 2023). In *Drosophila* embryos, loss of BAP1 leads to elevated H2AK119ub1 levels at Polycomb target genes and disrupts gene repression by impairing PRC1-mediated chromatin compaction (Bonnet et al., 2022). In mouse embryonic stem cells (mESCs), PR-DUB broadly promotes gene activation by limiting pervasive deposition of H2AK119ub1 across the genome, yet it is also responsible for Polycomb repression via a mechanism distinct from that in *Drosophila* (Conway et al., 2021; Fursova et al., 2021; Kolovos et al., 2020; Li et al., 2023). By contrast, studies in human cell lines have revealed that BAP1 functions as a transcriptional activator, acting to restrict PRC1-mediated H2AK119ub1 accumulation at gene regulatory elements, including enhancers (Campagne et al., 2019; Wang et al., 2018). These findings underscore the context-dependent role of BAP1 in gene regulation, suggesting that it may act to support both Polycomb repression and transcriptional activation, depending on cellular and developmental context. Thus, the precise mechanisms by which BAP1 regulates transcription remain enigmatic, and it is essential to dissect its *in vivo* function in balancing Polycomb-mediated silencing and transcriptional activation.

In this study, we aimed to identify the key H2AK119ub1 deubiquitinases (DUBs) in oocytes and early embryos, and to dissect how their activity contributes to the formation of facultative heterochromatin and euchromatin during oogenesis. We reveal that PR-DUB safeguards the oocyte epigenome and female fertility by protecting euchromatin from ectopic Polycomb activity, rather than by enforcing transcriptional repression.

## Results

### Identification of key H2AK119ub1 deubiquitinases in early embryos

To identify key DUBs responsible for H2AK119ub1 removal *in vivo*, we first analyzed the expression dynamics of known H2A DUBs during oogenesis and early embryonic development using publicly available RNA-seq and ribosome profiling datasets (Xiong et al., 2022; Zhang et al., 2022). Based on high expression levels, ribosome association in oocytes and preimplantation embryos, and lack of activity toward monoubiquitinated histone H2B, we selected five candidate H2A DUBs for further analysis: USP16 (Joo et al., 2007), USP28 (Li et al., 2019), USP21 (Nakagawa et al., 2008), BAP1 (Scheuermann *et al*., 2010), and MYSM1 (Zhu et al., 2007)(**Fig. s1a-d**). To assess their functional activity toward H2AK119ub1, we microinjected zygotes with Flag-tagged mRNAs encoding each candidate immediately after fertilization and evaluated H2AK119ub1 levels by immunostaining in late zygotes (∼6 hours post-injection) (**Fig. 1a**). Subcellular localization analyses revealed that USP16 and USP21 were primarily cytoplasmic, whereas BAP1, USP28, and MYSM1 exhibited no strong preference between nucleus and cytoplasm (**Fig. 1b**).

**Figure 1.**
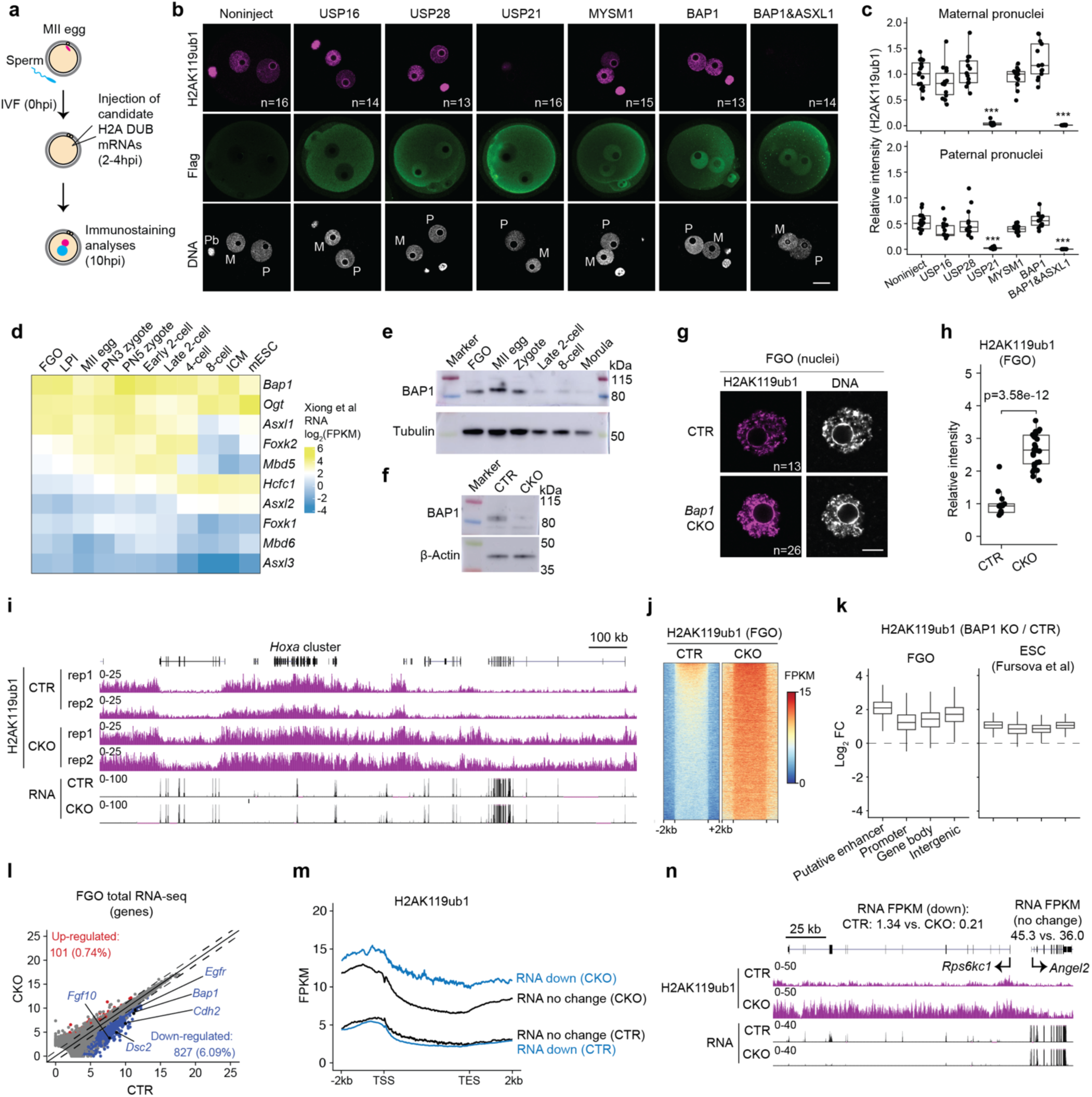
Loss of BAP1 causes pervasive increase of H2AK119ub1 and primarily results in gene downregulation in oocytes. a) Experimental design. MII: metaphase II; IVF: *in vitro* fertilization; DUB: deubiquitinase; hpi: hrs post IVF. **b)** Immunofluorescence images of H2AK119ub1 and Flag tag signals for non-injected zygotes, and zygotes injected with the indicated candidate H2A DUB mRNAs. Number of zygotes analyzed are indicated. M: maternal pronuclei; P: paternal pronuclei; Pb: polar body. Scale bar: 20 μm. **c)** Quantification of H2AK119ub1 fluorescence intensity from panel **(b)**. Boxplot: center line, median; box limits, 25th–75th percentiles; whiskers, ±1.5× interquartile range. ***: *p* < 0.001, two-sided Student’s *t*-test. **d)** RNA expression profiles of Polycomb Repressive Deubiquitinase (PR-DUB) subunits in mouse oocytes and preimplantation embryos. FGO: fully grown oocyte; LPI: late prometaphase I; PN: pronuclei; ICM: inner cell mass; mESC: mouse embryonic stem cell; FPKM: fragments per kilobase per million mapped reads. Data from public RNA-seq datasets (Xiong *et al*., 2022). **e)** Immunoblot showing dynamic expression of BAP1 across oocytes and preimplantation stages. 100 oocytes/embryos were used per sample for FGO, MII, zygote, and late 2-cell stages; 50 embryos were used for 8-cell and morula stages. **f)** Immunoblot confirming BAP1 depletion in FGOs from *Bap1* CKO mice. 100 oocytes per group. CTR: control; CKO: conditional knockout. **g)** Immunofluorescence images of H2AK119ub1 in nuclei of CTR and CKO FGOs. Number of analyzed FGOs are indicated. Scale bar: 10 μm. **h)** Quantification of H2AK119ub1 fluorescence intensity from panel **(g)**. Boxplot format as in **(c)**. *p*-value calculated by two-sided Student’s *t*-test. **i)** Genome browser views of H2AK119ub1 and RNA levels in FGOs at the indicate genomic region. **j)** Heatmaps of H2AK119ub1 levels at the H2AK119ub1 domains (see Methods, n=9,921) in CTR and CKO FGOs. **k)** Boxplot illustrating the changes of H2AK119ub1 at the indicated genomic regions in FGO and mESC. Boxplot format as in **(c)**. mESC H2AK119ub1 data from public datasets (Fursova *et al*., 2021). FC, fold change. **l)** Scatter plot comparing gene expression in CTR versus CKO FGOs. Red: upregulated; blue: downregulated genes in CKO. Differential expression criteria: fold change (FC) ≥ 2, adjusted *p* < 0.05, and FPKM ≥ 0.5. **m)** Metaplot showing H2AK119ub1 enrichment at gene loci that are unchanged or downregulated in *Bap1* CKO FGOs. TSS: transcription start site; TES: transcription ending sites. **n)** Genome browser views of H2AK119ub1 and RNA levels at the *Rps6kc1* and *Angel2* loci in FGOs.

Among these candidates, USP21 overexpression led to a robust depletion of H2AK119ub1 within 6 hours (**Fig. 1b, c**). In contrast, BAP1 alone did not affect H2AK119ub1 levels, but co-expression with ASXL1 significantly depleted the H2AK119ub1 signal. This finding aligns with previous studies showing that ASXL1 is required to activate BAP1’s H2A DUB function (Sahtoe et al., 2016). Overexpression of USP16, USP28, or MYSM1 had a mild impact, if any, on H2AK119ub1 levels, although we cannot exclude the possibility that they may function in a cell type– specific context or require additional cofactors. Together, these data suggest that USP21 and BAP1 potentially regulate H2AK119ub1 during early development. Since that the *Usp21* null mice are viable and fertile (Fan et al., 2014; Pannu et al., 2015), we selected BAP1 for detailed functional analysis in this study.

### BAP1 limits genome wide pervasive H2AK119ub1 accumulation in oocytes

Having identified BAP1 as a potential key H2A DUB in zygotes, we next examined the expression dynamics of other PR-DUB subunits during oogenesis and early embryonic development. Reanalysis of publicly available RNA-seq and ribosome profiling datasets (Xiong *et al*., 2022; Zhang *et al*., 2022) revealed that key PR-DUB components, including *Bap1, Asxl1, Foxk2,* and *Ogt*, are highly expressed and associated with ribosomes throughout oocyte maturation and preimplantation stages (**Fig. 1d, s1e**). Consistent with these findings, immunoblot analysis confirmed the presence of BAP1 protein in FGOs and early embryos (**Fig. 1e**).

To investigate the function of PR-DUB during oogenesis and early development, we generated an oocyte-specific conditional knockout (CKO) of *Bap1* by crossing a floxed *Bap1* line (see Methods), in which exons 6-12 are flanked by loxP sites, with a *Gdf9-iCre* transgenic line (Lan et al., 2004) (**Fig. s2a**). *Gdf9-iCre* is specifically expressed in oocytes of primordial follicles by postnatal day 3 (Lan *et al*., 2004), enabling targeted deletion of *Bap1* in early-stage oocytes. Cre-mediated recombination is predicted to produce a truncated BAP1 protein lacking catalytic activity and the ability to interact with ASXL partners (Sahtoe *et al*., 2016) (**Fig. s2b**). Throughout this study, *Bap1^fl/fl^*mice were used as controls (CTR), and *Gdf9-iCre; Bap1^fl/fl^* mice were used as CKO. At 6-9 weeks of age, comparable number of FGOs were recovered from CTR and CKO female mice (**Fig. s2c, d**). RNA-seq and immunoblot analyses confirmed effective depletion of *Bap1* at both the transcript and protein levels in CKO FGOs (**Fig. 1f, s2e**).

Immunofluorescence analyses revealed that H2AK119ub1 levels in CKO FGOs were significantly elevated, approximately 2- to 3-fold, compared to those in CTR oocytes (**Fig. 1g, h**). To determine the genomic regions where H2AK119ub1 increases, we performed spike-in normalized Cleavage Under Targets and Release Using Nuclease (CUT&RUN) (see Methods; **Fig. s2f**). Remarkably, this analysis uncovered a pervasive accumulation of H2AK119ub1 across the genome, with a relatively modest increase at actively transcribed genes (**Fig. 1i, j; Fig. s2g**). The extent of H2AK119ub1 elevation in BAP1-deficient oocytes appeared more pronounced than that in mESCs (Fursova *et al*., 2021) (**Fig. 1k**). Notably, putative enhancers, defined by distal H3K27ac peaks (Liu *et al*., 2024), exhibited a particularly strong increase in H2AK119ub1 (**Fig. 1k**), suggesting a role of BAP1 in regulating enhancer activities. Together, these findings indicate that BAP1 functions to restrain widespread H2AK119ub1 accumulation in oocytes.

### BAP1 primarily activates gene expression in oocytes

We next investigated transcriptomic changes resulting from *Bap1* deletion in FGOs. Comparative analysis identified 101 upregulated and 827 downregulated genes in CKO FGOs (**Fig. 1l, Table S1**). The predominance of downregulated genes suggests that BAP1 primarily acts as a gene activator in oocytes, likely by counteracting H2AK119ub1, a key player in PRC1-mediated gene silencing (Blackledge et al., 2020; Tamburri et al., 2020). Downregulated genes are enriched for gene ontology (GO) terms implicated in oogenesis, such as “cell adhesion molecule binding” (*Cdh2*, *Dsc2, Cdh18*) and “growth factor binding” (*Fgf10, Egfr*, *Foxp2*) (**Fig. s2h**). To link the observed transcriptional dysregulation to changes in H2AK119ub1, we examined H2AK119ub1 levels at these genes. While downregulated genes had similar baseline H2AK119ub1 enrichment in CTR oocytes, they acquired significantly higher levels of H2AK119ub1 in CKO FGOs compared to non-differentially expressed genes (DEGs) (**Fig. 1m, n**). Notably, this increase occurred across both genic and intergenic regions.

It has been previously reported that the methyl-binding protein MBD6 recruits PR-DUB to retrotransposons by recognizing m^5^C-modified chromatin-associated RNAs, a process critical for retrotransposon activation (Zou et al., 2024a). However, we found that BAP1 deficiency in oocytes had only a modest effect on the expression of repeat elements (**Fig. s2i, Table S1**). This suggests a limited role for MBD6-mediated PR-DUB recruitment in retrotransposon regulation during oogenesis, consistent with the low RNA-seq and Ribo-seq expression levels of *Mbd6* in oocytes (**Fig. 1d, s1e**). In sum, these findings indicate that the widespread accumulation of H2AK119ub1 following BAP1 loss primarily leads to gene repression, with a subset of genes being more susceptible than others (further discussed below).

### BAP1 preserves H3K27ac by preventing ectopic H3K27me3 deposition in oocytes

Previous studies have shown that H2AK119ub1 guides H3K27me3 deposition (Blackledge et al., 2014; Cooper et al., 2014; Kalb et al., 2014; Tavares et al., 2012), and that H3K27me3 and H3K27c are mutually exclusive. We therefore examined how BAP1 loss affects the H3K27me3 and H3K27ac landscapes in oocytes. Unlike the dramatic increase in H2AK119ub1, immunofluorescence analyses revealed that global levels of both H3K27me3 and H3K27ac remained largely unchanged in CKO FGOs (**Fig. 2a, b**). This observation was further supported by CUT&RUN analyses, which confirmed that most broad H3K27me3 and H3K27ac domains, known to be oocyte-specific (Liu *et al*., 2024; Zheng *et al*., 2016), were retained in CKO FGOs (**Fig. 2c, s3a**).

**Figure 2.**
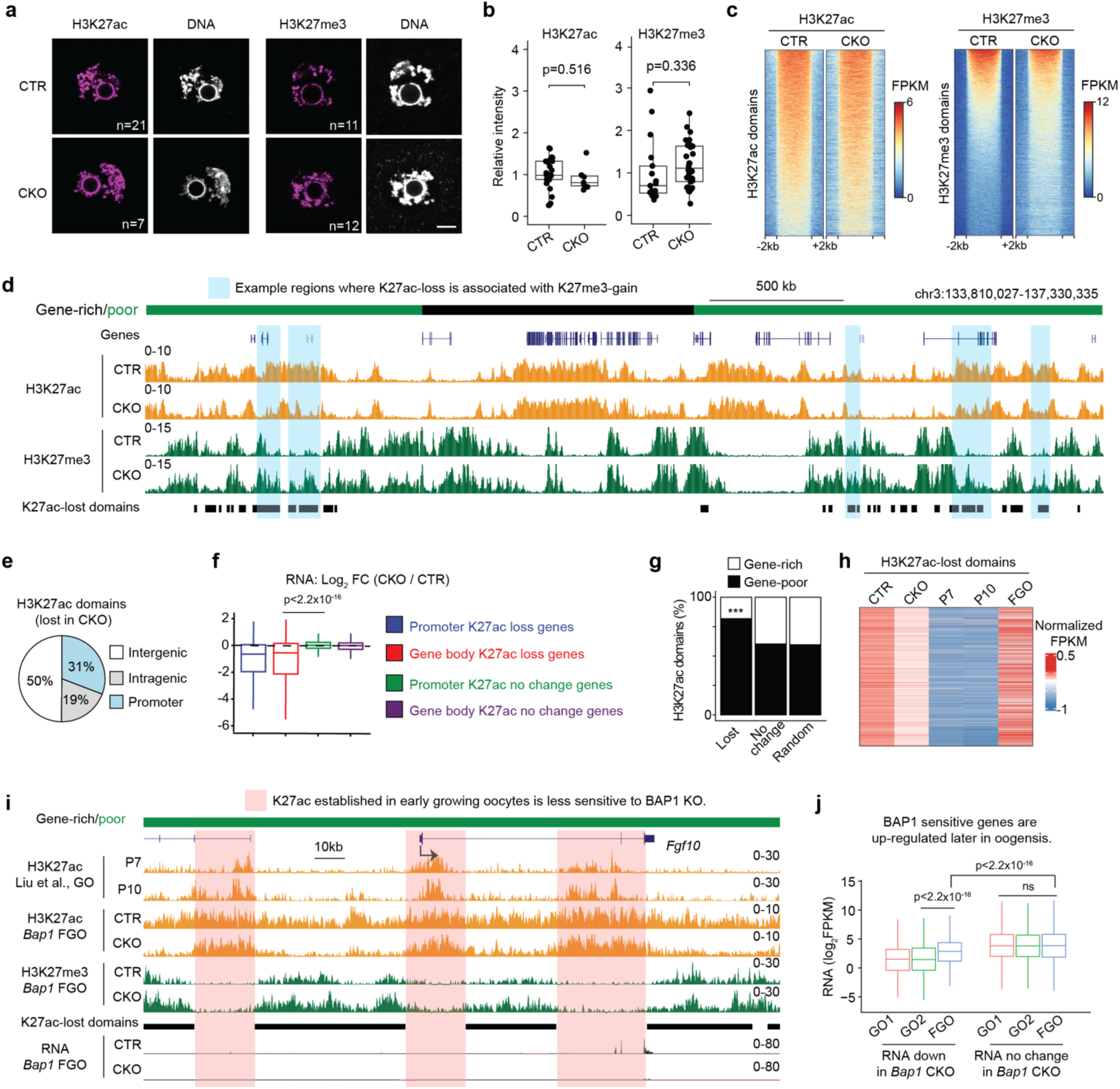
BAP1 preserves H3K27ac by preventing ectopic H3K27me3 deposition in oocytes. a) Immunofluorescence images of H3K27ac and H3K27me3 in nuclei of *Bap1* control (CTR) and conditional knockout (CKO) fully grown oocytes (FGOs). Number of FGOs analyzed are indicated. Scale bar: 10 μm. **b)** Quantification of H3K27ac and H3K27me3 intensities from panel **(a)**. Boxplot: center line, median; box limits, 25th–75th percentiles; whiskers, ±1.5× interquartile range. *p*-value: two-sided Student’s *t*-test. **c)** Heatmaps showing CUT&RUN signal intensity of H3K27ac and H3K27me3 across defined H3K27ac (n = 10,363) and H3K27me3 (n = 15,751) domains in *Bap1* CTR and CKO FGOs (see Methods). **d)** Genome browser view depicting H3K27ac and H3K27me3 signals at the indicated genomic locus. Regions with significantly reduced H3K27ac levels in CKO FGOs (“H3K27ac-lost domains”) are indicated. **e)** Genomic annotation of the H3K27ac-lost domains in *Bap1* CKO FGOs. **f)** RNA expression changes for genes associated with the indicated H3K27ac domain categories following *Bap1* deletion. Boxplot format as in **(b)**. *p*-value: two-sided Wilcoxon rank-sum test. **g)** Stacked bar plot showing the proportions of H3K27ac-lost and unchanged domains located in gene-rich versus gene-poor genomic regions. Chi-square test: \*\*\**p* < 2.2×10^-16^. **h)** Heatmap showing H3K27ac signal during oocyte growth across H3K27ac-lost domains. The H3K27ac data for postnatal day 7 (P7), P10 growing oocytes and FGOs are from public datasets (Liu *et al*., 2024). **i)** Genome browser view of H3K27ac and H3K27me3 profiles at the *Fgf10* locus. **j)** RNA expression levels of downregulated and unchanged genes upon *Bap1* depletion in wild-type growing oocytes (GO1 and GO2) and FGOs. GO1 and GO2 are oocytes from early secondary and secondary follicles, respectively. GO1/GO2/FGO RNA-seq are from public datasets (Zhang *et al*., 2020). Boxplot format as in **(b)**. *p*-value: two-sided Wilcoxon rank-sum test; ns: not significant.

However, closer inspection of genome browser tracks revealed specific loci where H3K27ac was reduced and H3K27me3 correspondingly increased in CKO FGOs (**Fig. 2d, s3b**). To systematically characterize these changes, we identified 9,043 H3K27ac-lost domains (ranging from 10 to 375 kb) in CKO oocytes (see Methods, **Table S2**). These domains already displayed relatively lower H3K27ac and higher H3K27me3 in CTR oocytes compared to regions with unchanged H3K27ac (**Fig. s3c**). H2AK119ub1 levels increased similarly in both H3K27ac-lost and H3K27ac-unchanged regions in CKO FGOs (**Fig. s3c**). Most H3K27ac-lost domains (69%) were in intergenic regions and gene bodies, suggesting potential enhancer activity (**Fig. 2e**). Genes associated with H3K27ac loss, either at promoters or within gene bodies, were preferentially downregulated in CKO FGOs (**Fig. 2f**). Together, these results indicate that BAP1 deficiency causes H3K27ac loss and H3K27me3 gain at selective loci, contributing to gene repression in oocytes.

To better understand the mechanisms underlying locus specific H3K27ac loss in CKO FGOs, we examined the genomic features of the affected regions. A striking characteristic of the H3K27ac-lost domains is their preferential localization within gene-poor regions, as opposed to gene-rich regions (**Fig. 2d, 2g, s3b**; see Methods). Previous studies have shown that H3K27ac peaks in gene-poor regions are more prevalent in FGOs compared to growing oocytes (Liu *et al*., 2024). Consistently, we found that these H3K27ac-lost domains exhibit low H3K27ac levels in growing oocytes at postnatal day 7 (P7) and day 10 (P10) but gain high levels of H3K27ac during the final stage of oocyte growth, as seen in FGOs (**Fig. 2h, 2i**). This suggests that regions unaffected by *Bap1* deletion acquire H3K27ac early during oocyte growth, whereas regions that lose H3K27ac in CKO oocytes gain it later and are therefore more vulnerable to *Gdf9-iCre*-mediated deletion of *Bap1* (**Fig. 2i, s3d**). Supporting this idea, only 9.9-13.4% of H3K27ac-lost domains overlapped with H3K27ac peaks in P7/P10 growing oocytes, whereas this overlap increased to 49-57.9% for H3K27ac-unchanged domains (**Fig. s3e**). Further supporting this model, genes unaffected by *Bap1* deletion are already highly expressed early during oocyte growth, whereas genes downregulated in CKO FGOs reach peak expression only at the fully grown stage and show lower expression levels than the other group (**Fig. 2j, s3f**). Together, these data suggest that the timing of H3K27ac establishment during oocyte development underlies the locus specific H3K27ac loss observed in *Bap1* CKO FGOs.

### BAP1 plays a limited role in Polycomb-mediated silencing in oocytes

Having established that BAP1 preserves oocyte-specific broad H3K27ac domains by preventing ectopic H3K27me3 deposition, we next examined its potential role in Polycomb-mediated gene silencing. In mESCs, it has been proposed that widespread accumulation of H2AK119ub1 in PR-DUB mutants may sequester PRC2, thereby diminishing H3K27me3 deposition at canonical Polycomb group (PcG) target loci (Conway *et al*., 2021; Fursova *et al*., 2021; Li *et al*., 2023). To evaluate this model in oocytes, we assessed H2AK119ub1, H3K27me3, and H3K27ac profiles at the promoters of typical PcG target genes as defined in mESCs (Matsuwaka et al., 2025). This analysis revealed a marked increase in H2AK119ub1 at PcG target promoters in *Bap1* CKO FGOs, accompanied by a relatively modest decrease in H3K27me3 and a corresponding gain in H3K27ac (**Fig. 3a, b, s4a**).

**Figure 3.**
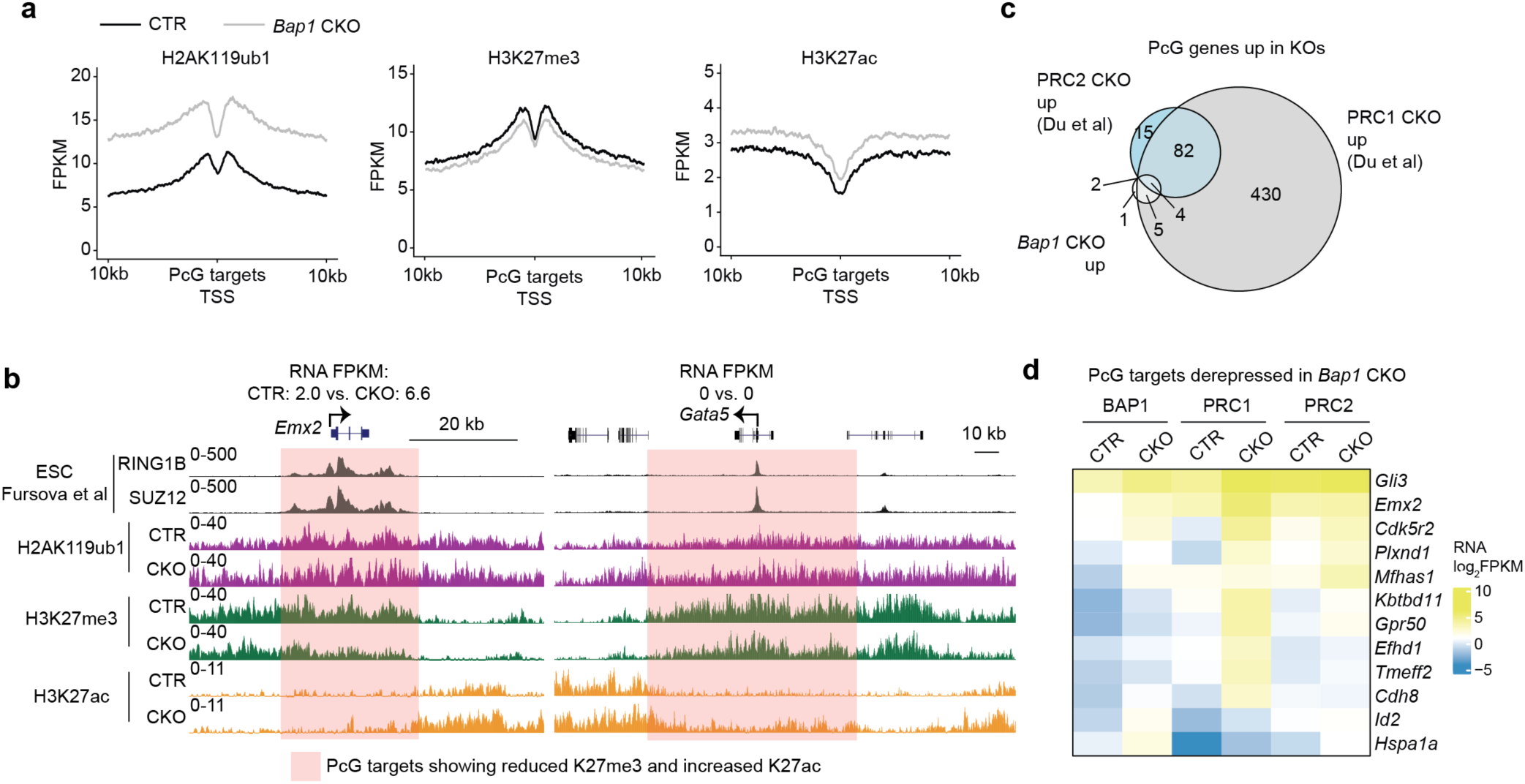
BAP1 plays a limited role in Polycom-mediated gene silencing in oocytes. a) Metaplots displaying signal intensities of H2AK119ub1, H3K27me3, and H3K27ac at Polycomb group (PcG) target genes in *Bap1* control (CTR) and conditional knockout (CKO) fully grown oocytes (FGOs) (see Methods). **b)** Genome browser views of H2AK119ub1, H3K27me3, and H3K27ac signals at the PcG target loci *Emx2* and *Gata5*. Public RING1B and SUZ12 ChIP-seq data in embryonic stem cells (ESCs) are included for comparison (Fursova et al., 2019). **c)** Venn diagram showing the overlap of PcG target genes that are de-repressed in PRC1, PRC2, and *Bap1* CKO FGOs. PRC1/2 RNA-seq are from public datasets (Du *et al*., 2020). **d)** Heatmap depicting expression levels of PcG targets that are de-repressed in *Bap1* CKO FGOs, compared across PRC1 and PRC2 CKO oocytes.

To assess whether the observed chromatin alterations could lead to transcriptional de-repression of PcG targets, we re-analyzed gene expression in *Bap1* CKO FGOs. For comparison, we examined publicly available RNA-seq datasets from PRC1 and PRC2 CKO oocytes (Du *et al*., 2020). Consistent with the established importance of PRC1 in postnatal oocyte development (Posfai *et al*., 2012), PRC1 loss led to a greater number of DEGs (2,417 upregulated and 700 downregulated) compared to PRC2 loss (414 upregulated and 67 downregulated) (**Fig. s4b, c**). Among these, 521 and 103 PcG targets were de-repressed in PRC1 and PRC2 CKO oocytes, respectively (**Fig. 3c**). By contrast, only 12 PcG targets were upregulated in *Bap1* CKO FGOs, and the extent of their upregulation was substantially lower than that observed in PRC1/2 mutants (**Fig. 3d**). These findings suggest that, although the global accumulation of H2AK119ub1 in BAP1-deficient oocytes may partially impair PRC2 recruitment and reduce H3K27me3 deposition at PcG targets, the overall integrity of Polycomb-mediated silencing is largely preserved. Taken together, these results indicate that BAP1 plays a limited role in the regulation of Polycomb-dependent gene silencing in oocytes.

### Loss of maternal BAP1 severely impairs preimplantation development

Having established the critical role of BAP1 in safeguarding the oocyte epigenome and transcriptome, we next assessed how maternal BAP1 deficiency may affect female fertility. To this end, CTR and CKO females were co-caged with wild-type males for six months. *Bap1* CKO females exhibited a significant reduction in fertility that was attributable to both fewer litters and smaller litter sizes (**Fig. 4a-c**). Ovarian morphologies of *Bap1* CKO females appear largely normal (**Fig. s5a**), which is consistent with the fact that comparable numbers of FGOs are retrieved from control and CKO females at 6-9 weeks of age (**Fig. s2c, d**), However, *in vitro* meiotic maturation analysis revealed a mild reduction in the metaphase II rate in CKO oocytes (**Fig. s5b, c**). Additionally, the number of embryos recovered at embryonic day 3.5 (E3.5) was slightly smaller in the CKO group (**Fig. 4d**), suggesting that ovulation is moderately impaired. Importantly, the embryos from *Bap1* CKO females showed defective preimplantation development, with most (∼90%) delayed at the 8-16-cell or morula stage at E3.5, whereas control embryos (∼96%) progressed to the blastocyst stage by this time point (**Fig. 4e, f**). Although extended *ex vivo* culture for an additional 24 hours allowed most mutant embryos to form blastocysts, these were of poor quality, exhibiting reduced total cell numbers and abnormal expression of lineage markers, including NANOG (epiblast), GATA4 (primitive endoderm), and CDX2 (trophectoderm) (**Fig. 4g, h**).

**Figure 4.**
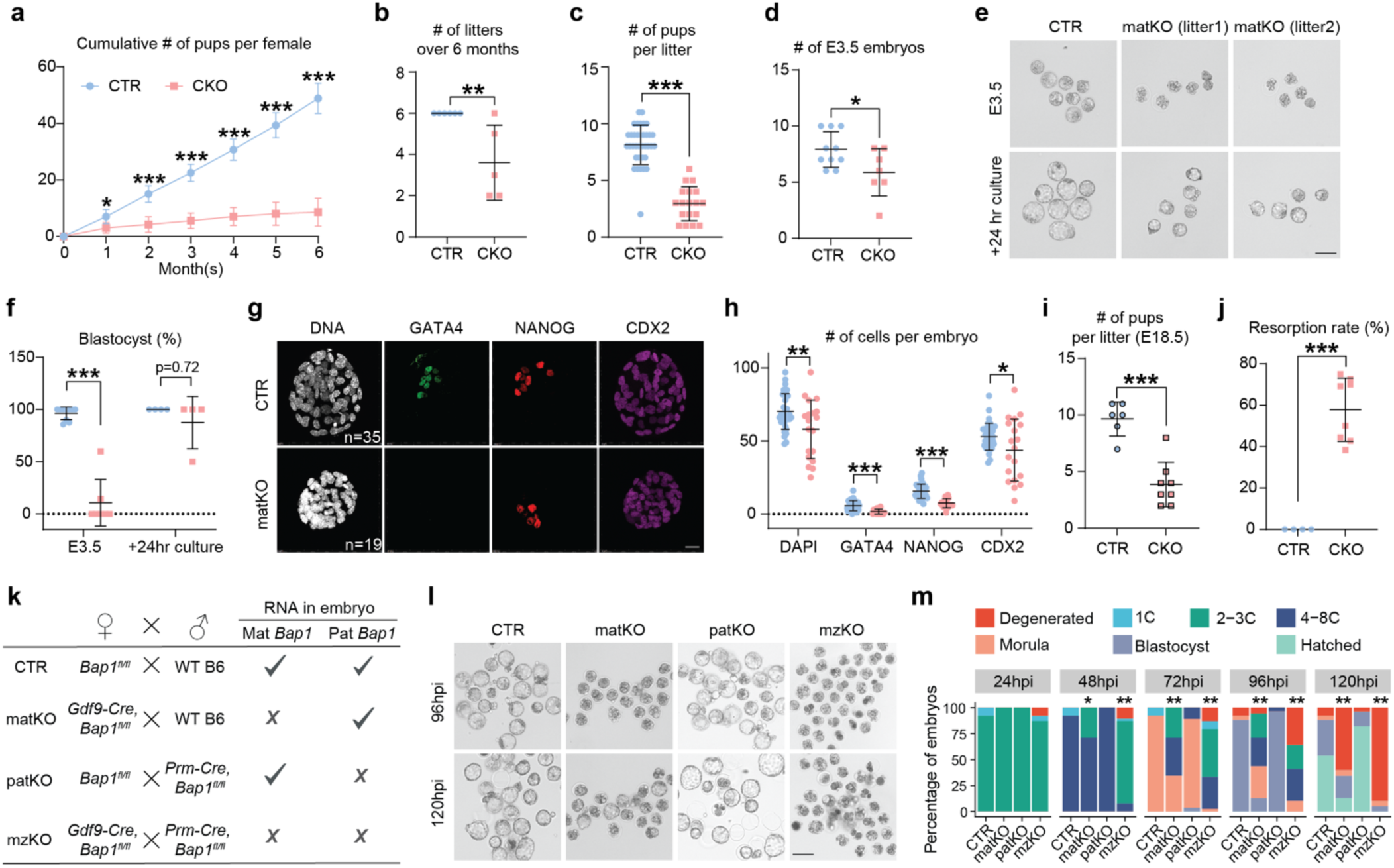
Loss of BAP1 severely impairs female reproductive capacity. a) Fertility assessment of *Bap1* control (CTR, *n* = 6) and conditional knockout (CKO, *n* = 5) female mice. Starting at 7 weeks of age, females were continuously housed with wild-type B6 males for six months. Data are presented as mean ± SD. \**p* < 0.05, \*\*\**p* < 0.001, two-sided Student’s *t*-test. **b)** Total number of litters produced during the 6-month mating trial for *Bap1* CTR (*n* = 6) and CKO (*n* = 5) females. Data are presented as mean ± SD. \*\**p* < 0.01, two-sided Student’s *t*-test. **c)** Average number of pups per litter. A total of 36 and 18 litters were analyzed for CTR and CKO groups, respectively. Data are presented as mean ± SD. \*\*\**p* < 0.001 two-sided Student’s *t*-test. **d)** Number of E3.5 embryos per litter from *Bap1* CTR (n=10) and CKO (n=7) females. Data are presented as mean ± SD. \**p*-values < 0.05, two-sided Student’s *t*-test. **e)** Brightfield images of E3.5 embryos collected from *Bap1* CTR and CKO females. Embryos were cultured *ex vivo* for an additional 24 hours. Scale bar, 100 μm. **f)** Percentages of CTR and maternal knockout (matKO) embryos that reached the blastocyst stage at E3.5 and after 24-hour culture. At E3.5, 79 embryos from 10 litters (CTR) and 41 embryos from 7 litters (matKO) were analyzed. For E3.5 + 24hr culture, 36 embryos from 4 litters (CTR) and 19 embryos from 4 litters (matKO) were analyzed. Data are presented as mean ± SD. Chi-square test: \*\*\**p* < 2.2×10^-16^. **g)** Immunofluorescence staining of CTR and matKO blastocysts (E3.5 + 24 hr culture) using antibodies against NANOG, GATA4, and CDX2. Number of blastocysts analyzed are indicated. Scale bar, 20 μm. **h)** Quantification of cell numbers for blastocysts shown in (**j**). Data are presented as mean ± SD. *p < 0.05, \*\**p* < 0.01, \*\*\**p* < 0.001, two-sided Student’s *t*-test. **i)** Number of viable pups per litter at embryonic day 18.5 (E18.5) in *Bap1* CTR (*n* = 6) and CKO (*n* = 8) groups. Data are presented as mean ± SD. \**p*-values < 0.001, two-sided Student’s *t*-test. **j)** Embryonic resorption rates at E18.5 for *Bap1* CTR (*n* = 6) and CKO (*n* = 8) females. Data are presented as mean ± SD. \*\*\**p*-values < 0.001, two-sided Student’s *t*-test. **k)** Schematics of IVF groups. Mat: maternal; Pat: paternal. **l)** Brightfield images of blastocysts of the indicated groups at 96 and 120 hrs post IVF (hpi). Scale bar, 100 μm. **m)** Stacked bar plot showing developmental progression of IVF embryos at specified time points. 26 (CTR), 55 (matKO), 28 (patKO), and 39 (mzKO) embryos were analyzed. 1C: one-cell stage. \**p* = 0.06, \*\**p* < 0.001, chi-square test.

To further evaluate developmental competence of the blastocysts, we performed cesarean section (C-section) at E18.5 and found that only 3.9 ± 2.0 matKO embryos reached term, representing just 41.4% of the number observed in CTR females (9.7 ± 1.5)(**Fig. 4i, j, s5d**), indicating a high rate of post-implantation embryonic loss likely attributable to compromised blastocyst quality. Notably, both placentae and fetuses from the maternal knockout (matKO) group were slightly heavier than those of controls (**Fig. s5e, f**), possibly due to reduced litter size. Thus, these data indicate that BAP1-deficient oocytes could not support normal preimplantation development to give rise to competent blastocysts.

We next sought to determine when developmental delays first arise in matKO embryos during preimplantation development. By monitoring embryos derived through *in vitro* fertilization (IVF), we observed that developmental delays began as early as the 2- to 4-cell transition (**Fig. 4k-m**). The delay became more pronounced by 72 hours post-IVF (hpi), at which point only ∼30% of CKO embryos had reached the morula stage. Ultimately, only ∼30% of CKO embryos developed into blastocysts, in contrast to the ∼85% blastocyst rate observed in the control group. The more pronounced developmental arrest observed in IVF-derived matKO embryos likely reflects the increased sensitivity of mutant embryos to suboptimal *ex vivo* culture conditions relative to the *in vivo* environment.

Because the paternal *Bap1* allele remains intact in matKO embryos (**Fig. 4k**), it is possible that *Bap1* expression from the paternal genome during zygotic genome activation (ZGA) at the 2-cell stage partially compensates for the maternal loss. To test this hypothesis, we generated *Prm-Cre; Bap1^fl/fl^* male mice, in which *Bap1* is deleted during the late haploid spermatid stage. When *Bap1* CKO oocytes were fertilized with sperm from these males, the resulting embryos, referred to as maternal-zygotic knockout (mzKO), displayed markedly more severe developmental defects. Few mzKO embryos progressed beyond the 2- to 4-cell stages. The observed phenotype was not due to intrinsic defects in *Prm-Cre; Bap1^fl/f^* sperm, as their fertilization of control oocytes resulted in normal progression to the blastocyst stage. Together, these findings demonstrate that maternal BAP1 is essential for preimplantation development, and loss of both maternal and zygotic *Bap1* further exacerbates the phenotype.

### Loss of maternal BAP1 causes maternal-to-zygotic transition (MZT) defects

Having established that maternal BAP1 is essential for preimplantation development, we next sought to investigate the underlying mechanisms responsible for the developmental arrest and/or delay observed in *Bap1* matKO embryos. The MZT is a critical developmental window during which, maternal RNAs and proteins are degraded, and embryonic development becomes increasingly dependent on zygotic gene expression initiated during ZGA (Kojima et al., 2025). Minor ZGA occurs at late 1-cell (L1C) and early 2-cell (E2C) stages with the activation of hundreds of genes, whereas major ZGA takes place at late 2-cell (L2C) stage, during which thousands of genes are transcriptionally activated (Kravchenko and Tachibana, 2025; Zou et al., 2024b).

Given that the developmental defect in *Bap1* matKO embryos emerges as early as the 2- to 4-cell transition, we hypothesized that loss of maternal BAP1 disrupts the MZT. To test this hypothesis, we performed total RNA sequencing of control and *Bap1* mutant oocytes/embryos at the MII stage, L1C, E2C, and L2C stages (**Fig. 5a, s6a**). Comparative transcriptome analyses identified 886, 869, 1096, and 2740 DEGs between CTR and mutant groups at the MII stage, L1C, E2C, and L2C stages, respectively (**Fig. 5b, Table S1**). The number of DEGs was comparable across the GV, MII, and L1C stages, with a predominant trend of gene downregulation in *Bap1* mutants (**Fig. 1i and 5b**). Notably, ∼67% (511 genes) of L1C-downregulated genes were already repressed in mutant oocytes (**Fig. 5c**), and these genes were enriched for GO terms related to cell adhesion (e.g., *Cdh2*, *Cdh18*, *Cdh20*, *Dsc2*) and growth factor binding (e.g., *Gata3*, *Fgf10*, *Egfr*, *Foxp2*) (**Fig. s6b**). Thus, these data suggest the aberrant transcriptomes in *Bap1* CKO oocytes are inherited into 1-cell embryos.

**Figure 5.**
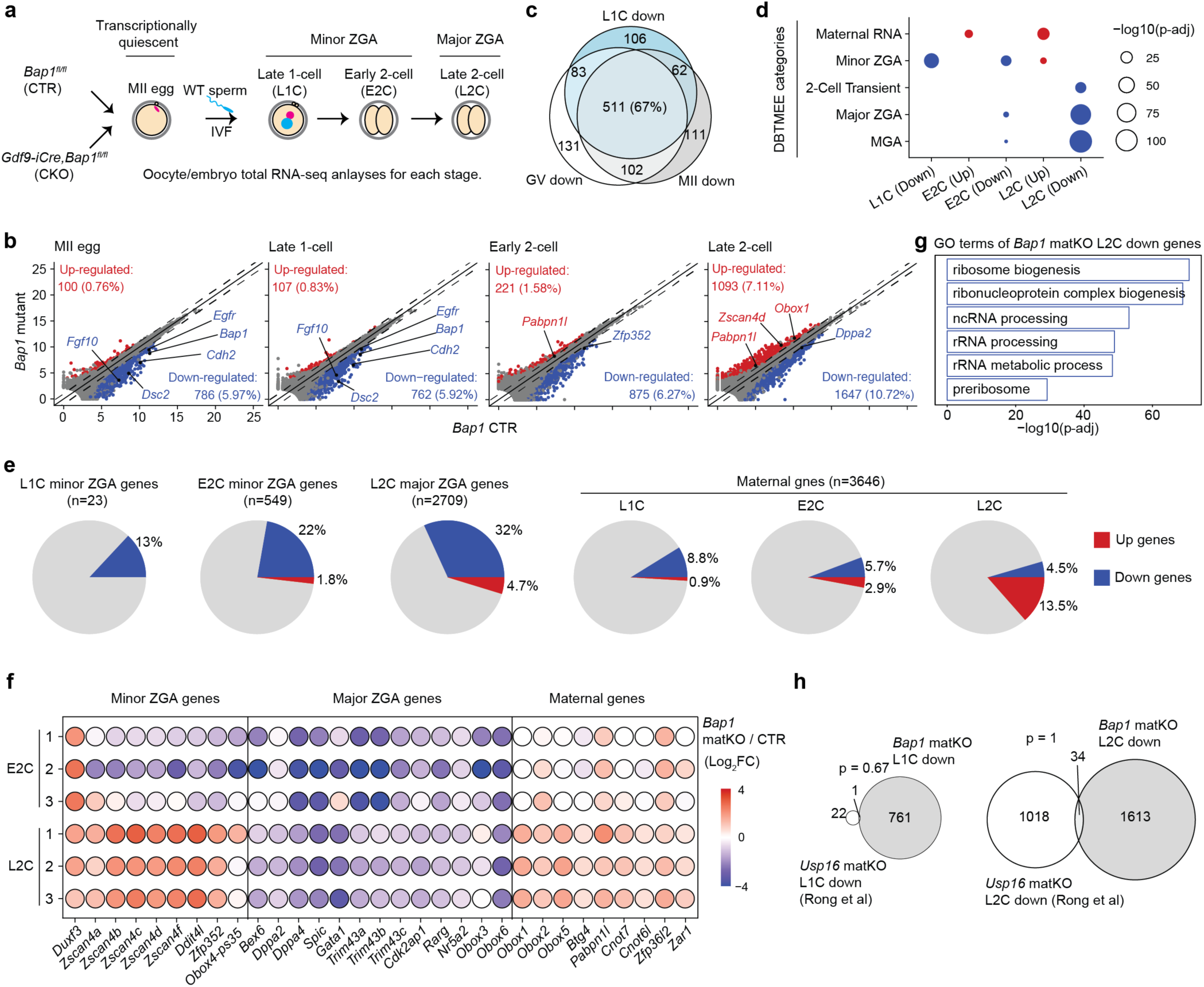
Loss of maternal BAP1 causes defective maternal-to-zygotic transition. a) Schematic of sample collection timeline for total RNA-seq analysis. ZGA: zygotic genome activation; MII: metaphase II eggs. **b)** Scatter plots comparing gene expression between *Bap1* control (CTR) and conditional knockout (CKO) oocytes/embryos at the indicated developmental stages. Red and blue dots indicate significantly upregulated and downregulated genes in *Bap1* mutant, respectively. Differential expression was defined by fold change ≥ 2, adjusted *p* < 0.05, and FPKM ≥ 0.5. **c)** Venn diagram illustrating the overlap of downregulated genes in *Bap1* mutant samples at GV, MII, and L1C stages. **d)** Bubble plot showing enrichment of differentially expressed genes in selected gene categories from the DBTMEE database. Statistical significance determined using a hypergeometric test; adjusted *p*-values are indicated. **e)** Pie charts showing the proportion of downregulated ZGA genes and upregulated maternal genes in *Bap1* matKO embryos. Gene categories were defined according to Wang et al., 2022. **f)** Balloon plot depicting RNA expression dynamics of representative maternal, minor ZGA, and major ZGA genes at E2C and L2C stages. Three biological replicates were analyzed for each condition. **g)** GO terms enriched among downregulated genes in *Bap1* matKO L2C embryos. **h)** Venn diagrams illustrating the overlap of differentially expressed genes between *Bap1* and *Usp16* maternal KO embryos. RNA-seq data for *Usp16* samples are from public datasets (Rong *et al*., 2022). *p*-value: hypergeometric test.

We next analyzed compositions of DEGs at E2C and L2C stages. Downregulated genes at these stages significantly overlapped with those classified as “minor ZGA,” “major ZGA,” and “2-cell transient” in the DBTMEE v2 database (Park et al., 2015), while upregulated genes were enriched for transcripts categorized as “maternal RNA” (**Fig. 5d**), implicating defects in both zygotic activation and maternal RNA decay. To further assess the impact of maternal BAP1 loss on ZGA, we compared DEGs in *Bap1* mutants with previously defined zygote and E2C minor ZGA gene sets (Wang *et al*., 2022). Among 23 zygote minor ZGA genes detected in our dataset, three (*Sox6*, *Stk39*, *Lcp2*) were significantly downregulated in *Bap1* matKO embryos at L1C (**Fig. 5e**), with additional minor ZGA genes such as *Dux* and *Usp17la/c/d* showing a trend of downregulation (**Table S1**). Furthermore, 22% of minor ZGA genes and 32% of major ZGA genes were downregulated at the E2C and L2C stages, respectively (**Fig. 5e, f**), including key regulators of early development such as *Zscan4*, *Cdk2ap1*, *Nr5a2*, and *Obox3/6* (**Fig. 5f**)(Falco et al., 2007; Festuccia et al., 2024; Gassler et al., 2022; Guo et al., 2024; Ji et al., 2023; Lai et al., 2023; Modzelewski et al., 2021; Zhao et al., 2024b). Downregulated genes at L2C were also significantly enriched for GO terms related to ribosome biogenesis and rRNA processing (**Fig. 5g**), potentially accounting for the developmental defects at the 2- to 4-cell transition. In addition, *MT2/MERVL* retrotransposons—known to be essential for early development (de la Rosa et al., 2024; Sakashita et al., 2023; Yang et al., 2024)—were markedly downregulated in *Bap1* mutants at the E2C stage (**Fig. s6c**). Interestingly, a subset of E2C minor ZGA genes showed aberrant upregulation at L2C, possibly reflecting delayed downregulation (**Fig. 5f**). Lastly, 13.5% of maternal transcripts were upregulated at the L2C stage, indicating defective maternal RNA degradation (**Fig. 5e**). Collectively, these findings demonstrate that loss of maternal BAP1 causes defects in both ZGA and maternal transcript clearance during MZT.

Ubiquitin-specific peptidase 16 (USP16) has previously been implicated in H2A deubiquitination during the GV-to-MII transition and is essential for mouse ZGA (Rong et al., 2022). To evaluate whether USP16 and BAP1 act through similar or distinct mechanisms during oogenesis and early embryogenesis, we reanalyzed the published transcriptomic data. To our surprise, there was minimal overlap between DEGs in *Usp16* and *Bap1* mutants at both L1C and L2C stages (**Fig. 5h, s6d**), suggesting that maternal USP16 and BAP1 function via distinct regulatory pathways. Consistent with this notion, loss of BAP1, but not USP16, causes increase in H2AK119ub1 in FGOs (**Fig. 1)**(Rong *et al*., 2022). These findings underscore the mechanistic divergence between USP16 and BAP1 in regulating H2A deubiquitination during development and highlight the need for further investigation into their individual and potentially complementary roles.

### Maternal BAP1 regulates embryonic enhancer activity during ZGA

Having demonstrated MZT defects in *Bap1* matKO embryos, we next investigated how the loss of maternal BAP1 affects chromatin dynamics during early embryogenesis. To this end, we performed immunostaining for H2AK119ub1, H3K27me3, and H3K27ac in CTR and *Bap1* matKO embryos at the 1-cell (6 and 10 hpi), L2C (28 hpi), and 4-cell (48 hpi) stages. Since developmental arrest in *Bap1* matKO embryos becomes evident as early as the 2- to 4-cell transition, only morphologically normal 4-cell embryos were included in the analysis. Similar to FGOs, H2AK119ub1 levels were significantly elevated in *Bap1* matKO embryos compared to CTR at all stages examined (**Fig. 6a, s7**). In contrast, H3K27me3 immunostaining levels were largely comparable between the two groups. Interestingly, H3K27ac signals were markedly reduced at the 2-cell and 4-cell stages in the absence of maternal BAP1 (**Fig. 6a, s7**), suggesting impaired formation of active euchromatin that likely occurs independently of H3K27me3.

**Figure 6.**
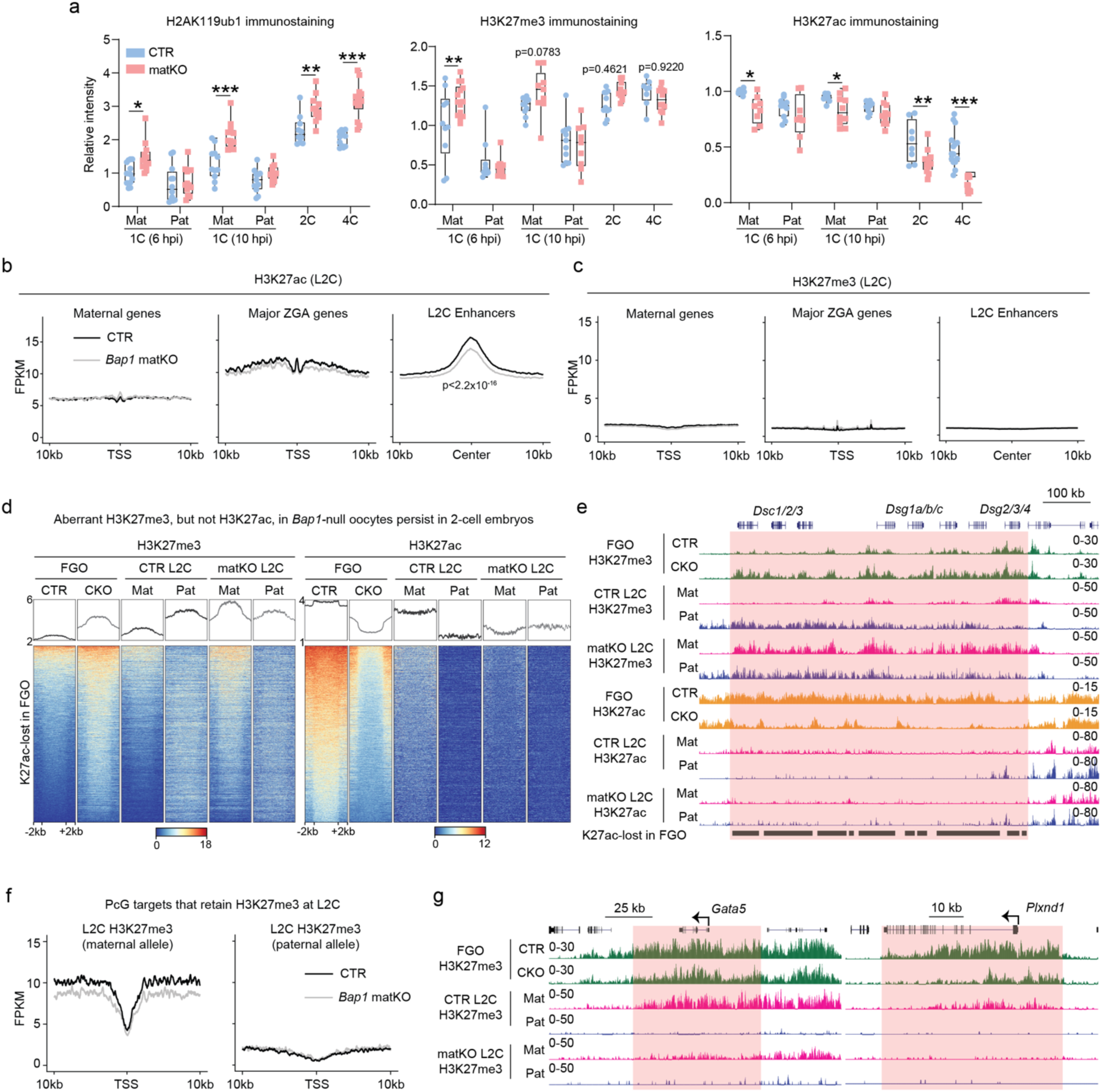
Aberrant H3K27me3 landscapes in *Bap1* null oocytes, but not H3K27ac changes, persist in early embryos. a) Quantification of H2AK119ub1, H3K27me3, and H3K27ac fluorescence intensities. Boxplot: center line, median; box limits, 25th–75th percentiles; whiskers, ±1.5× interquartile range. * *p* < 0.05, **: *p* < 0.01, ***: *p* < 0.001, two-sided Student’s *t*-test. Mat: maternal pronuclei; Pat: paternal pronuclei; hpi: hrs post IVF. **b)** Metaplot showing H3K27ac enrichment at maternal genes, major ZGA genes, and enhancers in *Bap1* CTR and matKO late 2-cell (L2C) embryos. TSS: transcription start site. *p*-value: two-sided Wilcoxon rank-sum test. **c)** Metaplot showing H3K27me3 enrichment same as in panel **b)**. **d)** Metaplots (top) and heatmaps (bottom) showing changes in H3K27me3 and H3K27ac profiles at the H3K27ac-lost domains in fully grown oocytes (FGOs) and L2C embryos from *Bap1* CTR and CKO groups. For L2C embryos, allelic signals are shown separately for maternal (Mat) and paternal (Pat) alleles (see Methods). **e)** Genome browser views of H3K27me3 and H3K27ac profiles in FGOs and L2C embryos at the indicated genomic loci. **f)** Metaplots showing allelic H3K27me3 enrichment at a subset of Polycomb group (PcG) targets in late 2-cell embryos from *Bap1* CTR and matKO groups. **g)** Genome browser views of H3K27me3 profiles in FGOs and L2C embryos at the indicated genomic loci.

To further explore this, we performed CUT&RUN profiling of H3K27ac and H3K27me3 at the L2C stage, when major ZGA occurs. Consistent with the immunofluorescence data, the global H3K27me3 landscape was largely unchanged between control and matKO embryos, whereas H3K27ac signals were moderately decreased (**Fig. s8a, b**). Notably, putative L2C enhancers, defined by distal H3K27ac peaks (Liu et al., 2024), were strongly reduced in matKO embryos (**Fig. 6b, s8c**). In contrast, H3K27ac levels at promoter regions of both maternal and major ZGA genes remained largely unaffected by the loss of maternal BAP1 (**Fig. 6b**). These findings suggest that impaired activation of L2C enhancers contributes to defective ZGA in maternal BAP1-deficient embryos. Moreover, H3K27ac levels at the embryonic enhancers were low in FGOs and showed no difference between control and mutant oocytes (**Fig. s8d**), indicating that this defect arises post-fertilization. Furthermore, H3K27me3 levels at maternal genes, major ZGA loci, and L2C enhancers were comparable between control and matKO embryos (**Fig. 6c**), excluding aberrant H3K27me3 deposition as a contributing factor. Collectively, these results indicate that maternal BAP1 is required for proper embryonic enhancer activities during ZGA, independent of H3K27me3 regulation.

### Aberrant H3K27me3 landscape, but not H3K27ac changes, in BAP1-null oocytes persists into early embryos

Given that H3K27me3-deficient states in *Eed*- or *Pcgf1/6*-null oocytes are irreversibly transmitted to early embryos (Inoue *et al*., 2018; Mei *et al*., 2021), we next examined whether the aberrant H3K27me3 landscape observed in *Bap1*-null FGOs (**Fig. 2–3**) similarly persists post-fertilization. Leveraging the fact that our L2C embryos are derived from B6/129 × PWK F_1_ hybrids, we performed allele-specific analysis of H3K27me3 distribution. We first focused on genomic regions, primarily located in gene deserts, that acquire ectopic H3K27me3 and corresponding H3K27ac loss in *Bap1*-deficient oocytes (**Fig. 2**). In control L2C embryos, these regions displayed paternally biased H3K27me3 enrichment (**Fig. 6d, e and s8e**), consistent with prior reports that paternal H3K27me3 in preimplantation embryos predominantly localizes to gene-poor regions (Zheng *et al*., 2016). Remarkably, these loci acquired maternal-specific H3K27me3 enrichment in *Bap1* matKO embryos, and this ectopic gain was already evident in CKO FGOs (**Fig. 6d, e and s8e**), suggesting inheritance of the abnormal chromatin state from the oocyte. In contrast, H3K27ac levels at these loci declined sharply from FGO to L2C stages in both control and mutant embryos, with the CTR maternal allele showing only a slightly higher H3K27ac than that in mutants (**Fig. 6d, e and s8e**). This suggests that this H3K27ac reduction reflects a normal developmental progression and is not specific to BAP1 loss. Notably, the inherited ectopic H3K27me3 may not initiate gene repression in embryos, as the downregulated genes within these domains were already transcriptionally reduced at earlier stages (**Fig. s8f, g**), implying that these transcriptional defects arise during oogenesis rather than being induced post-fertilization.

Lastly, we assessed H3K27me3 and H3K27ac profiles at the typical Polycomb targets in control and *Bap1* matKO L2C embryos. Previous studies have shown that H3K27me3 is largely lost from promoter regions of Polycomb targets following fertilization, with only a subset retaining residual H3K27me3 during preimplantation development (Matsuwaka *et al*., 2025; Zheng *et al*., 2016). Analysis of the Polycomb targets that retain H3K27me3 at the L2C stage (n = 719; see Methods) revealed a maternal bias in H3K27me3 enrichment in control embryos (**Fig. 6f, g**). In contrast, this maternal H3K27me3 signal was either completely lost or markedly reduced in *Bap1* matKO embryos (**Fig. 6f, g**). Notably, the reduction observed in mutant L2C embryos was more pronounced than that in *Bap1* CKO oocytes, suggesting progressive loss post-fertilization. By contrast, the modest H3K27ac gain seen at Polycomb targets in CKO FGOs was not sustained in early embryos (**Fig. s8h, i**). Together, these results demonstrate that the aberrant H3K27me3 landscape, but not H3K27ac changes, established in BAP1-null oocytes is inherited into early embryonic stages.

## Discussion

In this study, we report that BAP1 is essential for establishing spatially distinct chromatin domains during oogenesis, specifically active euchromatin marked by H3K27ac and facultative heterochromatin marked by H2AK119ub1 and H3K27me3. BAP1 prevents pervasive accumulation of H2AK119ub1 across the genome and protects broad, oocyte-specific H3K27ac domains, particularly in gene-poor regions, from ectopic H3K27me3 deposition. This chromatin regulation is critical for activating a subset of oocyte-specific genes, including those involved in cell adhesion and growth factor signaling. Consequently, loss of maternal BAP1 leads to impaired developmental competence of oocytes and female subfertility. Notably, our findings demonstrate that BAP1 functions primarily as a transcriptional activator in oocytes, rather than enhancing Polycomb-mediated gene silencing. Together, these results reveal an essential role for PR-DUB in safeguarding the oocyte epigenome and preserving reproductive potential.

This study, together with previous work in *Drosophila* embryos (Bonnet *et al*., 2022), human cell lines (Campagne *et al*., 2019; Wang *et al*., 2018), and mESCs (Conway *et al*., 2021; Fursova *et al*., 2021; Kolovos *et al*., 2020; Li *et al*., 2023), underscores a conserved role for BAP1 in limiting widespread H2AK119ub1 accumulation, albeit to varying degrees across model systems. However, BAP1-null oocytes uniquely exhibit a striking loss of H3K27ac and a corresponding gain of H3K27me3 over broad genomic domains, often spanning hundreds of kilobases (**Fig. 7a**). Two non-mutually exclusive mechanisms may explain how increased H2AK119ub1 leads to H3K27me3 gain and H3K27ac loss in *Bap1* CKO oocytes. First, elevated H2AK119ub1 may enhance recruitment of PRC2.2 (Blackledge and Klose, 2021), resulting in deposition of H3K27me3, which in turn could displace H3K27ac and repress transcription. Supporting this, PRC2.2 depletion prevents intergenic H3K27me3 accumulation in *Bap1*-null mESCs (Conway *et al*., 2021). Second, ASXLs—components of PR-DUB—interact with the MLL3/4-containing COMPASS complex, which includes the H3K27 demethylase KDM6A (Szczepanski et al., 2020; Wang *et al*., 2018; Zhang et al., 2024; Zhao et al., 2022). In addition, BAP1-dependent enhancer binding of KDM6A has been observed in human cell lines (Wang *et al*., 2018). Thus, loss of BAP1 may therefore impair COMPASS targeting to enhancers, leading to increased H3K27me3 and reduced H3K27ac. Further investigation is needed to distinguish between these mechanisms and to elucidate how PR-DUB integrates with other chromatin-modifying complexes during oogenesis.

**Figure 7.**
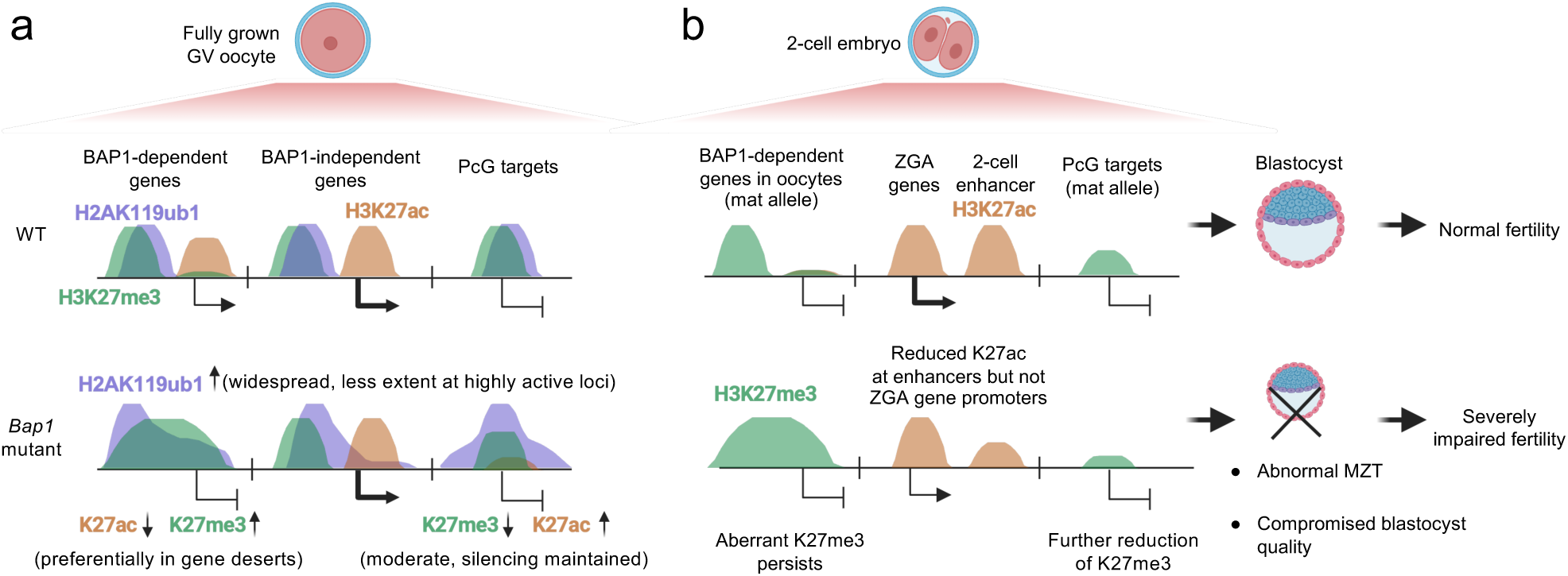
A model illustrating the functions of PR-DUB in oocyte epigenome and female fertility. a) BAP1 depletion causes pervasive increase of H2AK119ub1 in fully grown oocytes (FGOs), with a less extent at highly transcribed gene loci. Genes within gene deserts are preferentially down-regulated in *Bap1-*null FGOs, which is associated with increased H3K27me3 and reduced H3K27ac. Polycomb silencing is largely maintained at the target genes in FGOs despite moderate reduction of H3K27me3 and increase of H3K27ac. **b)** The aberrant H3K27me3 landscapes, but not H3K27ac, established in *Bap1-*null oocytes persist in early embryos. H3K27ac at enhancers, but not at ZGA gene promoters, are reduced in *Bap1* maternal KO embryos. The abnormal maternal-to-zygotic transition (MZT) in *Bap1* maternal KO embryos impairs preimplantation development, leading to compromised blastocyst quality and female subfertility.

It is unexpected that H3K27ac loss and H3K27me3 gain in BAP1-null oocytes preferentially occur in gene-poor regions, a phenomenon not observed in other model systems. Our findings suggest that the timing of H3K27ac establishment during oocyte development underlies this gene desert–biased loss of H3K27ac in *Bap1* CKO FGOs. Specifically, gene-poor regions acquire high levels of H3K27ac during the final stages of oocyte growth (Liu *et al*., 2024) and are therefore more sensitive to *Bap1* deletion than regions that gain H3K27ac earlier. Supporting this notion, *Bap1-*dependent genes reach peak expression only at the fully grown stage, whereas genes unaffected by *Bap1* deletion are already highly expressed early during oocyte growth. These observations suggest that in non-dividing cells like growing oocytes, BAP1 is particularly important for the establishment of new H3K27ac marks and activation of transcription, rather than for the maintenance of existing chromatin and transcriptional states. A less likely scenario is that PR-DUB is preferentially recruited to gene-poor regions via an unknown mechanism. To further dissect these possibilities, it would be informative to delete *Bap1* earlier than what is achieved with *Gdf9-iCre*–mediated CKO, to assess whether early BAP1 loss affects both gene-dense and gene-poor regions similarly.

Another unique aspect of BAP1 function in oocytes is its predominant role in promoting gene activation, with only a minor contribution to Polycomb-mediated gene silencing (**Fig. 7a**). This contrasts with findings in mESCs, where BAP1 loss leads to pervasive accumulation of H2AK119ub1, resulting in displacement of PRC1 and PRC2 from Polycomb target genes and consequent gene derepression (Conway *et al*., 2021; Fursova *et al*., 2021; Li *et al*., 2023). In BAP1-null oocytes, although H3K27me3 levels at canonical PcG targets are moderately reduced, this reduction appears insufficient to trigger derepression of these genes. One possible explanation is that the high abundance of PRC1 and PRC2 complexes accumulated during oocyte growth may buffer against sequestration by ectopic H2AK119ub1. Even if a portion of PRC1/2 is mis-localized, the remaining complexes might still be sufficient to maintain transcriptional repression at their target sites.

Interestingly, in *Drosophila*, PR-DUB preserves Polycomb silencing through a distinct mechanism compared to mESCs. In flies, H2AK118ub1 is rapidly removed by PR-DUB at embryonic stages when Polycomb repression is active and thus does not co-localize with H3K27me3 (Bonnet *et al*., 2022). In the absence of PR-DUB, elevated H2AK118ub1 disrupts chromatin compaction, impairing gene silencing (Bonnet *et al*., 2022). In this context, H2AK118ub1 may function as a transcriptional activator. A similar activating role for H2AK119ub1 has also been proposed in mammalian systems (Li et al., 2025; Zhao et al., 2024a). However, we did not observe such a scenario in mouse oocytes, suggesting that this mechanism is either not conserved in oocytes or that H2AK119ub1 increase in this context does not reproduce the chromatin environment seen in other systems.

The role of BAP1 in chromatin regulation is critical for the activation of a subset of oocyte-specific genes during oogenesis, particularly those involved in cell adhesion and growth factor signaling. Failure to activate these genes compromises the developmental competence of *Bap1*-null oocytes, and matKO embryos fail to properly complete the MZT, resulting in defective preimplantation development. Among the BAP1-dependent genes, *Desmocollin 3* (*Dsc3*) has been shown to be essential for preimplantation development (Den et al., 2006). Additionally, *Fibroblast growth factor 10* (*Fgf10*) has been reported to enhance oocyte maturation and developmental competence in bovine (Zhang et al., 2010). Other candidates that may contribute to the compromised oocyte and early embryonic development include *N-cadherin* (*Cdh2*) and *Epidermal growth factor receptor* (*Egfr*). These findings suggest that PR-DUB function during oogenesis is essential for supporting early embryonic development. However, we cannot exclude the possibility that maternal BAP1 also directly regulates embryonic enhancer activity after fertilization, as indicated by reduced H3K27ac at L2C enhancers in *Bap1* matKO embryos—a defect not observed in mutant FGOs.

The locus-specific gain of H3K27me3 in *Bap1*-null oocytes provides key insights into intergenerational epigenetic inheritance. In previous studies, H3K27me3-deficient states in *Eed*- or *Pcgf1/6*-null oocytes were shown to be irreversibly transmitted to early embryos (Inoue *et al*., 2018; Mei *et al*., 2021). Notably, *Pcgf1/6* matKO embryos fail to re-establish H3K27me3 domains lost in the oocyte, despite having functional PRC2 complexes (Mei *et al*., 2021). In this study, we observed that the aberrant H3K27me3 landscapes in *Bap1-*null oocytes, including ectopic gain in gene-poor regions and reduction at PcG targets, persists in early embryos (**Fig. 7b**). In contrast, the abnormal H3K27ac distributions in *Bap1-*deficient oocytes is largely reversed along with the global reduction of H3K27ac levels from FGO to L2C transition. It will be of interest in future studies to determine whether intergenerational inheritance is specific to heterochromatin-associated marks such as H3K27me3, but not euchromatin marks like H3K27ac, in mammals. The functional consequences of the inherited ectopic H3K27me3 in L2C embryos remain unclear. The downregulated genes within these regions at L2C are already transcriptionally repressed at earlier stages, suggesting that the gene expression defects originate during oogenesis rather than being induced post-fertilization. Nevertheless, it will be important to investigate how long these aberrant H3K27me3 domains persist and whether they contribute to gene regulation at later developmental stages.

## Materials & Methods

### Mouse breeding

To generate oocyte specific *Bap1* KO, *Bap1^flox^*animals (Jax strain #: 031874) were crossed with *Gdf9-iCre* transgenic mice (Jax strain #: 011062) to produce *Gdf9-iCre, Bap1 ^flox/+^* males and *Bap1 ^flox/+^* females. These mice were intercrossed to obtain *Bap1 ^flox/flox^*and *Gdf9-iCre, Bap1 ^flox/flox^*females, which were used as CTR and CKO, respectively. To generate sperm specific *Bap1* KO, *Bap1^flox^* animals were crossed with *Prm-Cre* mice (Jax strain #: 007252). The resulting *Prm-Cre, Bap1 ^flox/+^* females were subsequently bred with *Bap1 ^flox/+^*males to obtain *Prm-Cre, Bap1 ^flox/flox^*males for downstream experiments. Genotypes were confirmed by PCR using genomic DNA extracted from tail biopsies. Genotyping primers are listed in Table S3. All animal experiments were performed in accordance with the guidelines of the Institutional Animal Care and Use Committee at Cincinnati Children’s Hospital Medical Center.

### Mouse superovulation, IVF, embryo culture, and fertility assessment

To induce superovulation, 8-12-week-old female mice were intraperitoneally injected with either 7.5 IU Pregnant Mare Serum Gonadotropin (PMSG; BioVendor) or ∼0.15 mL CARD HyperOva (Cosmo Bio). After 44-48 hours, 7.5 IU Human Chorionic Gonadotropin (hCG; Sigma) was administered via intraperitoneal injection. Approximately 16-18 hours post-hCG injection, cumulus-oocyte complexes (COCs) were collected from the oviductal ampullae. IVF was performed using sperm isolated from the cauda epididymis of 9-15-week-old male mice. For collection of *Bap1* CTR and matKO embryos used in immunostaining, RNA-seq, and developmental rate analyses, sperm were obtained from wild-type C57BL/6J (Jax strain #: 000664) or *Prm-Cre, Bap1^flox/flox^*males and capacitated in Human Tubal Fluid (HTF; Millipore Sigma) medium for ∼1 hour at 37 °C in a CO₂ incubator with ambient air. For CUT&RUN experiments, sperm were isolated from wild-type PWK/PhJ mice (Jax strain #: 003715) and capacitated in CARD FERTIUP Preincubation Medium (Cosmo Bio).

The time of sperm addition to COCs was defined as 0 hours post-insemination (hpi). At ∼6 hpi, zygotes exhibiting two pronuclei were selected, washed in M2 medium (Millipore), and cultured in KSOM medium (Millipore) at 37 °C in a CO₂ incubator under ambient air conditions. Brightfield images of oocytes and embryos were acquired using EVOSII cell imaging system (Thermo Fisher Scientific). Collection of FGOs, MII oocytes for immunostaining, RNA-seq, and CUT&RUN experiments was described previously (Chen et al., 2025). *In vitro* maturation analysis for FGOs was described previously (Zhang et al., 2020).

For fertility assessment and embryo collection at E3.5 and E18.5, *Bap1* CTR and CKO females were co-caged with wild-type C57BL/6J males (Jax strain #: 000664). The presence of a vaginal plug the following morning was designated as E0.5. For E3.5 collections, blastocysts were flushed from the uterus using M2 medium (Millipore). For E18.5 collections, pregnant females were euthanized, and the number and weight of fetuses and placentas were assessed.

### Whole-mount immunostaining and histology analyses

Immunostaining was described previously (Inoue *et al*., 2017). Primary antibodies were rabbit anti-H3K27me3 (1:500, Cell Signaling, 9733), rabbit anti-H2AK119ub1 (1:2,000, Cell Signaling, 8240), rabbit anti-H3K27ac (1:500, Millipore Sigma, 07360), mouse anti-Flag (1:200, Sigma, F1804), rabbit anti-NANOG (1:200, Cosmo Bio, ATL-HPA072117-25), Goat anti-CDX2 (1:200, R&D, AF3665), or Mouse anti-GATA4 (1:200, R&D MAB2606). Secondary antibodies were Alexa Fluor 488 donkey anti-mouse IgG (1:200), Alexa Fluor 568 donkey anti-rabbit IgG (1:200), or Alexa Fluor 647 donkey anti-goat IgG (1:200) (Life Technologies). Fluorescence was detected under a laser-scanning confocal microscope (Nikon A1R inverted), and images were analyzed using ImageJ (NIH). For histology analyses, paraffin-embedded ovaries were sectioned at 5 μm, and stained with hematoxylin and eosin.

### Plasmid construction and mRNA preparation

Coding sequences for mouse *Usp16* (NM_024258), *Usp21* (NM_013919), *Usp28* (NM_175482), *Mysm1* (NM_177239), *Bap1* (NM_027088), and the N-terminal region (amino acids 1–476) of *Asxl1* (NM_015338) were synthesized by Twist Bioscience. Each sequence was cloned into the pcDNA3.1-Flag-poly(A)83 vector (Inoue and Zhang, 2014), inserted between an N-terminal Flag tag and an 83-nucleotide polyadenylation sequence. Plasmids were linearized by restriction enzyme digestion and used as templates for *in vitro* transcription with the mMESSAGE mMACHINE® T7 Ultra Kit (Life Technologies), following the manufacturer’s instructions. The resulting mRNA was purified by LiCl precipitation and quantified using a NanoDrop spectrophotometer.

### Micro-injection of H2A DUB mRNAs

IVF experiments using BDF1 (Jax strain #: 100006) MII oocytes and sperm were the same as in previous sections. At 2 hpi, fertilized zygotes with the presence of 2^nd^ polar body or fertilization cone were transferred into M2 medium (Sigma), and mRNA was injected into the cytoplasm of zygotes (∼2–4 hpi) using a Piezo impact-driven micromanipulator (Eppendorf). Following injection, embryos were cultured in HTF (Millipore) medium for an additional 4 h before being transferred to KSOM (Millipore). The mRNA concentrations for each H2A DUB mRNA were 2 µg µl^−1^ except that 1 µg µl^−1^ were used for *Bap1* and *Asxl1* mRNAs when they were co-injected.

### Western blotting

Oocytes and embryos collection and immunoblot analysis were as previously described (Zhang *et al*., 2020). Primary antibodies against BAP1 (1/1000 dilution; CST13271), Tubulin (1/1000 dilution; CST2144), Beta-actin (1/1000, sc-47778) were used in this study.

### Total RNA-seq and CUT&RUN library preparation and sequencing

RNA-seq libraries were generated using the SMART-Seq Stranded Kit (Takara) following the manufacturer’s instructions. CUT&RUN library preparation was described previously (Chen *et al*., 2025). For H2AK119ub1 CUT&RUN, *Drosophila* S2 nuclei were used for spik-in normalization. The number of S2 nuclei used for each experiment are included in Table S3. The libraries were sequenced on the Illumina NovaSeq X Plus platform at Novogene (2 × 150bp). The number of oocytes and embryos used for RNA-seq/CUT&RUN, sequencing depth, and reads alignment are included in Table S3.

### Total RNA-seq data processing

Adaptor trimming, read alignment, and read count generation for both genes and repeat subfamilies were performed as previously described (Yang *et al*., 2024). Differential gene expression analysis was carried out using DESeq2 (v1.38.3)(Love et al., 2014), applying the criteria: *adjusted p-value < 0.05* and *fold change ≥ 2*. For a gene to be considered differentially expressed, the higher-expression group was additionally required to have *FPKM > 0.5*. Gene Ontology (GO) term enrichment among differentially expressed genes (DEGs) was assessed using the enrichGO function in the clusterProfiler R package (v4.6.2)(Xu et al., 2024). BigWig files for visualization were generated using deepTools (v2.0.0)(Ramirez et al., 2016) with the following parameters: --binSize 25 -- normalizeUsing RPKM --scaleFactor 1.

Gene sets corresponding to zygote minor ZGA genes, early 2-cell minor ZGA genes, late 2-cell major ZGA genes, and maternal transcripts were obtained from (Wang *et al*., 2022). Enrichment of DEGs across gene categories defined in the DBTMEE_v2 database (Park *et al*., 2015) was analyzed using the enrichr function in clusterProfiler (v4.6.2)(Xu *et al*., 2024).

### CUT&RUN data processing

Raw paired-end reads from CUT&RUN experiments were trimmed using TrimGalore (v0.6.6) (https://github.com/FelixKrueger/TrimGalore) with the --paired option. For FGOs, trimmed reads were aligned to the mm10 mouse reference genome using Bowtie2 (v2.4.2)(Langmead and Salzberg, 2012) with the parameters: --no-unal --no-mixed --no-discordant -I 0 -X 1000. Uniquely mapped reads were filtered using Sambamba (v0.6.8) (Tarasov et al., 2015) with the criteria: -F “mapping_quality >= 30 and not (unmapped or mate_is_unmapped)”. PCR duplicates were removed using Picard (v2.18.22) (http://broadinstitute.github.io/picard/). BigWig coverage files were generated using deepTools (v2.0.0)(Ramirez *et al*., 2016) with the options: --binSize 25 --normalizeUsing RPKM --scaleFactor 1.

For late 2-cell (L2C) stage CUT&RUN data, to avoid alignment bias toward the reference genome (C57BL/6), reads were aligned to a custom genome where single-nucleotide polymorphisms (SNPs) between PWK/PhJ and B6/129 strains were masked with “N”. SNPsplit (v0.3.2)(Krueger and Andrews, 2016) was used to determine parental origin of read pairs. For H2AK119ub1 CUT&RUN datasets, spike-in normalization was performed by aligning reads to the *Drosophila melanogaster* reference genome (fly46), and scaling factors were calculated as described by (Skene and Henikoff, 2017)(see Table S3).

Reproducibility between biological replicates was assessed by calculating the Pearson correlation coefficient of FPKM values computed in non-overlapping 5-kb genomic bins. For downstream analyses, biological replicates were pooled using the merge command from Sambamba (v0.6.8)(Tarasov *et al*., 2015). The heatmaps of CUT&RUN profiles were generated using deepTools (v2.0.0)(Ramirez *et al*., 2016) computeMatrix and plotHeatmap.

### Identification of H3K27ac-lost domains in *Bap1* CKO FGOs

FPKM values were computed in non-overlapping 5-kb genomic bins across the genome. Bins with FPKM ≥ 2 and fold change ≥ 1.5 between conditions were classified as differentially enriched bins. To define broader enriched regions, adjacent differentially enriched 5-kb bins were merged using BEDTools (v2.30.0) (Quinlan and Hall, 2010). Only merged regions larger than 10 kb were retained and defined as differentially enriched domains. The domains were annotated using ‘‘makeTxDbFromGFF’’ and ‘‘annotatePeak’’ function from the ‘‘ChIPpeakAnno’’, ‘‘GenomicFeatures’’, and ‘‘ChIPseeker’’ Bioconductor R packages (Lawrence et al., 2013; Yu et al., 2015; Zhu et al., 2010).

### Analyses at PcG targets

The list of PcG target genes was obtained from (Matsuwaka *et al*., 2025). To identify PcG targets that retain H3K27me3 at the late 2-cell stage, FPKM values at promoter regions (±2.5 kb around the transcription start site) were calculated. Genes with FPKM ≥ 2 were considered as retaining H3K27me3. This cutoff was determined based on visual inspection of representative loci, and the resulting gene list was found to be comparable to (Matsuwaka *et al*., 2025).

### Identification H2AK119ub1, H3K27me3 and H3K27ac domains

A hidden Markov model (HMM)-based domain caller (Chen *et al*., 2021) was used to identify broad domains in control FGO and late 2-cell embryos.

### Annotations and genomic intervals

The gene annotation file (GFF3 format) was downloaded from GENCODE (release M25) (https://www.gencodegenes.org/). Gene promoters were defined as ±2.5 kb surrounding the transcription start sites (TSS). Gene bodies were defined from the TSS to the annotated transcription end site (TES). Intergenic regions were defined as genomic regions that do not overlap with annotated promoters or gene bodies. Putative enhancers specific to FGOs and late 2-cell embryos were obtained from (Liu *et al*., 2024). Gene-rich and gene-poor genomic regions were defined as previously described (Liu et al., 2018). All mm9-based genomic coordinates were converted to the mm10 assembly using the liftOver function from the rtracklayer R package (v1.58.0)(Lawrence et al., 2009).

### Visualization and statistical analyses

All statistical analyses were performed using R (v4.2.2). Statistical tests and sample sizes are indicated in the figure legends. All genomic browser tracks were visualized using the UCSC genome browser (Kent et al., 2002). Figure 7 was generated using BioRender.

### Public datasets used in this study

All public ChIP-seq, CUT&RUN, and RNA-seq datasets were processed using pipelines and parameters similar to those used for in-house data. Relevant datasets are cited in the figure legends and main text whenever applicable.

## Acknowledgements

We thank Dr. Yueh-Chiang Hu for rederiving the *Gdf9-iCre* mice. We are grateful to Amanda Barbosa and Dr. Matthew Kofron for assistance with imaging analysis. We also thank Drs. S.K. Dey, Tony De Falco, Xiaofei Sun, and Maria Mikedis for helpful discussion. Confocal microscopy was performed at the CCHMC Bio-Imaging and Analysis Facility. This work was supported by Z.C.’s start-up funds and the Eunice Kennedy Shriver National Institute of Child Health and Human Development (R00 HD104902). Z.C. is also supported by R01 HD118540.

## Author contributions

Z.C. conceived the project. Z.C. and J.K. designed the experiments. J.K. and Z.C. performed the experiments and analyzed the data, with assistance from P.L., S.I., and L.C. Z.C. carried out the bioinformatic analyses. Z.C. and J.K. wrote the manuscript with input from all authors.

## Data availability

All sequencing data were deposited in the Gene Expression Omnibus under accession number GSE302804 (token for reviewers:) and GSE302844 (token for reviewers:).

## Code availability

The code for this study is available at https://github.com/ZYChen-lab/Bap1_in_oogenesis_story.

## Declaration of generative AI in Scientific Writing

During the preparation of this work, the authors used ChatGPT to make the final manuscript more concise. After using this tool, the authors reviewed and edited the content as needed and take full responsibility for the content of the publication.

## Supplementary tables

Table S1. Differential gene and repeat expression analyses in *Bap1* CTR and mutant oocytes and embryos.

Table S2. List of H3K27ac-lost domains in *Bap1* CKO FGOs.

Table S3. List of datasets generated in this study and oligos used.

---

**Figure S1.**
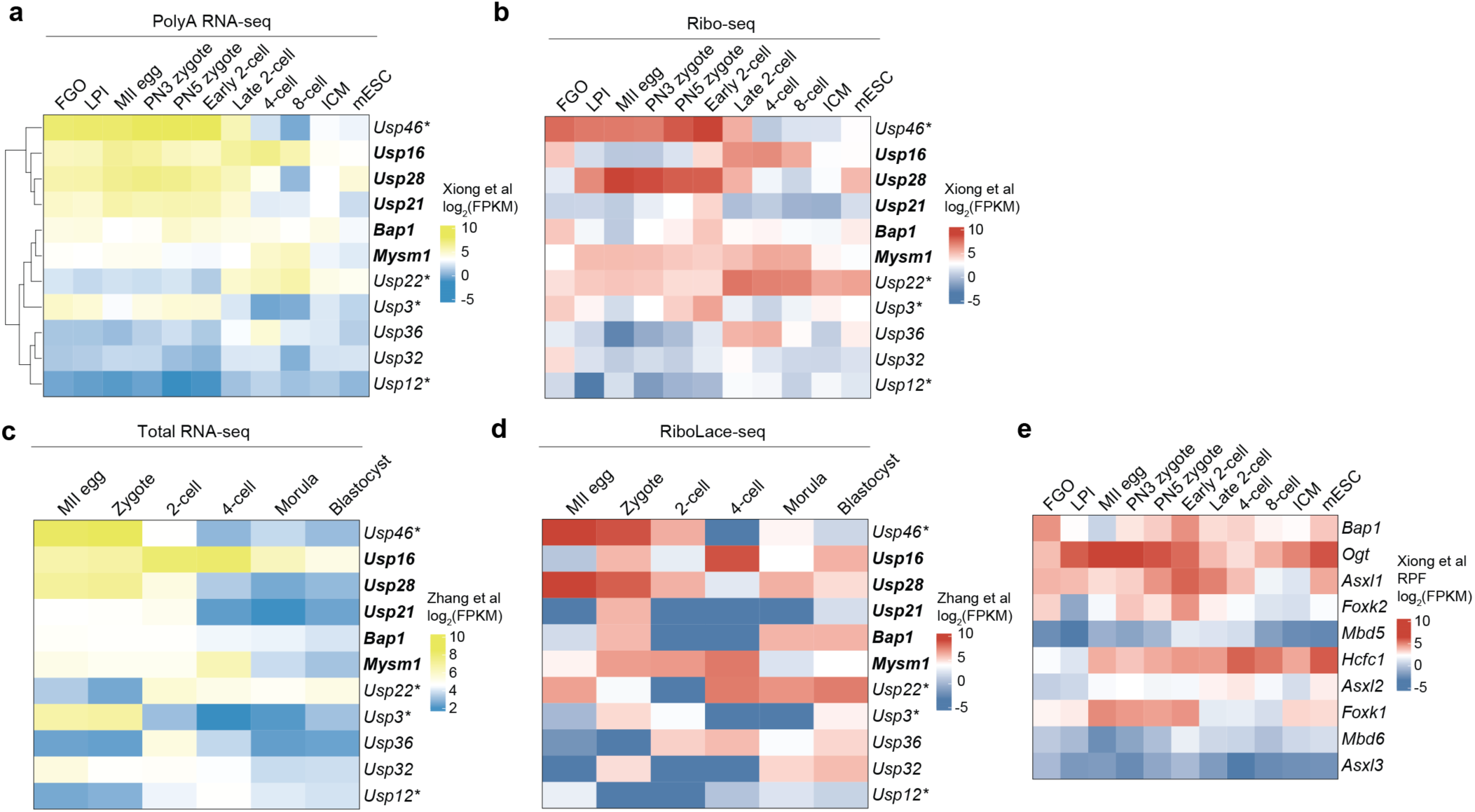
RNA sequencing (RNA-seq) and Ribosome profiling (Ribo-seq) signals for known H2A DUBs (a-d), and PR-DUB complex subunits (e). FGO: fully grown oocyte; LPI: late prometaphase I; MII: metaphase II; PN: pronuclear stage; ICM: inner cell mass; mESC: mouse embryonic stem cells. Expression values are presented as FPKM (fragments per kilobase of transcript per million mapped reads). * DUBs that also catalyzes H2B monoubiquitin. Data from public Ribo-seq datasets (Xiong *et al*., 2022; Zhang *et al*., 2022).

**Figure S2.**
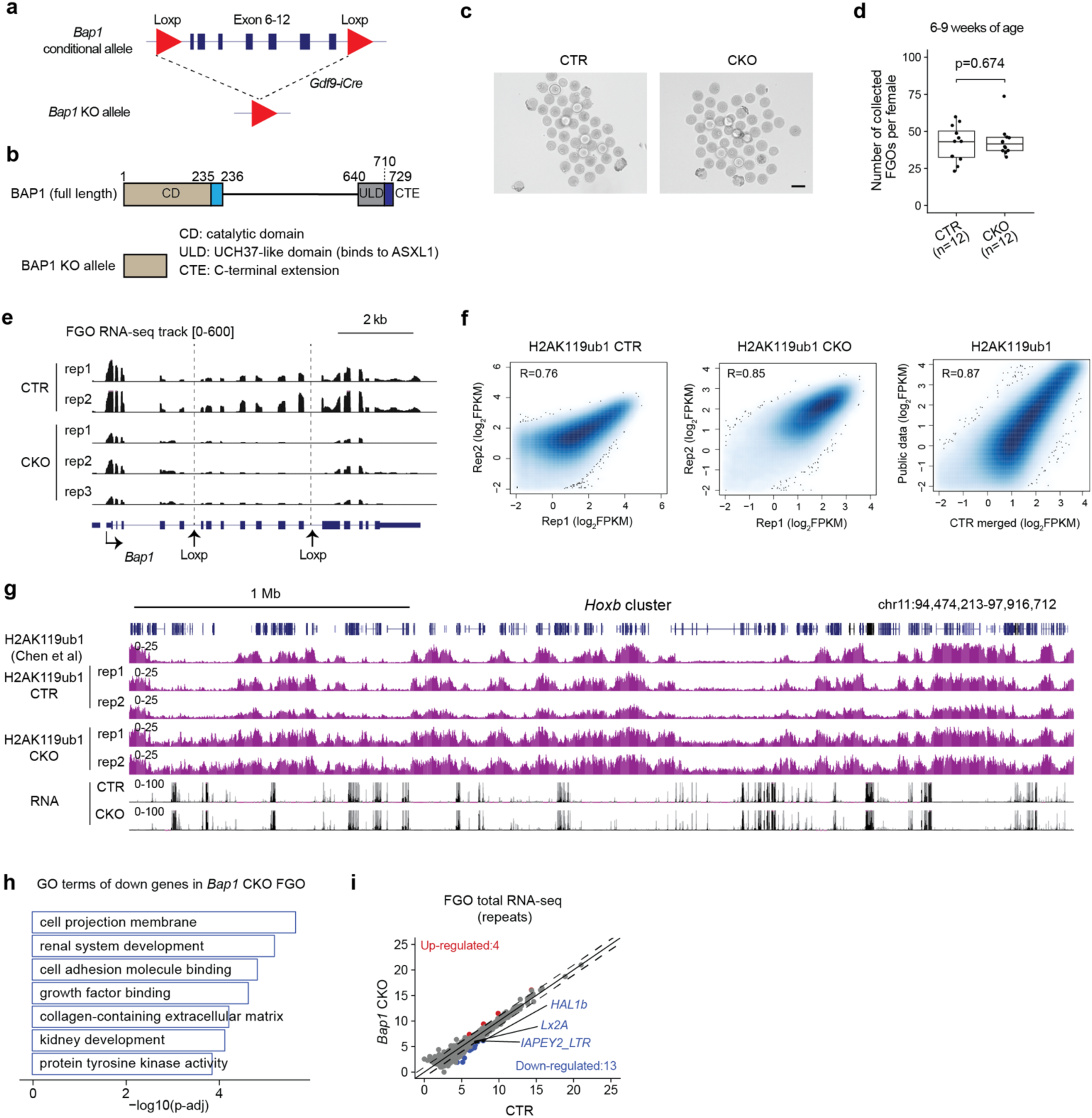
Generation and characterization of the *Bap1* conditional knockout (CKO) mouse model. a) Schematic showing the generation of *Bap1* CKO mice. **b)** Domain structures of full-length and truncated BAP1 proteins in the knockout model. **c)** Brightfield images of fully grown oocytes (FGOs) collected from one female mouse (6–9weeks old) per group. Scale bar, 100 μm. **d)** Boxplot showing the number of FGOs retrieved per female mouse in *Bap1* CTR and CKO groups. Sample sizes are as indicated. Boxplot features: center line indicates the median; box bounds represent the 25th and 75th percentiles; whiskers extend to ±1.5× the interquartile range. *p* value: two-sided Student’s *t*-test. **e)** Genome browser views showing RNA-seq signal tracks at the *Bap1* locus in CTR and CKO FGOs. **f)** Scatter plots showing reproducibility of H2AK119ub1 CUT&RUN profiles between biological replicates in FGOs. Correlation between *Bap1* CTR and a published dataset (Chen *et al*., 2021) isshown. Pearson correlation coefficients are indicated. **g)** Genome browser views of H2AK119ub1 CUT&RUN and RNA-seq signal tracks the indicated genomic locus in FGOs. **h)** Gene ontology (GO) terms enriched for the downregulated genes in *Bap1* CKO FGOs. **i)** Scatter plot comparing differential expression of repeat element subfamilies in *Bap1* CTR versus CKO FGOs. Red and blue dots indicate significantly upregulated and downregulated repeats in *Bap1* CKO, respectively. Differential expression was defined by fold change ≥ 2 and adjusted *p* < 0.05.

**Figure S3.**
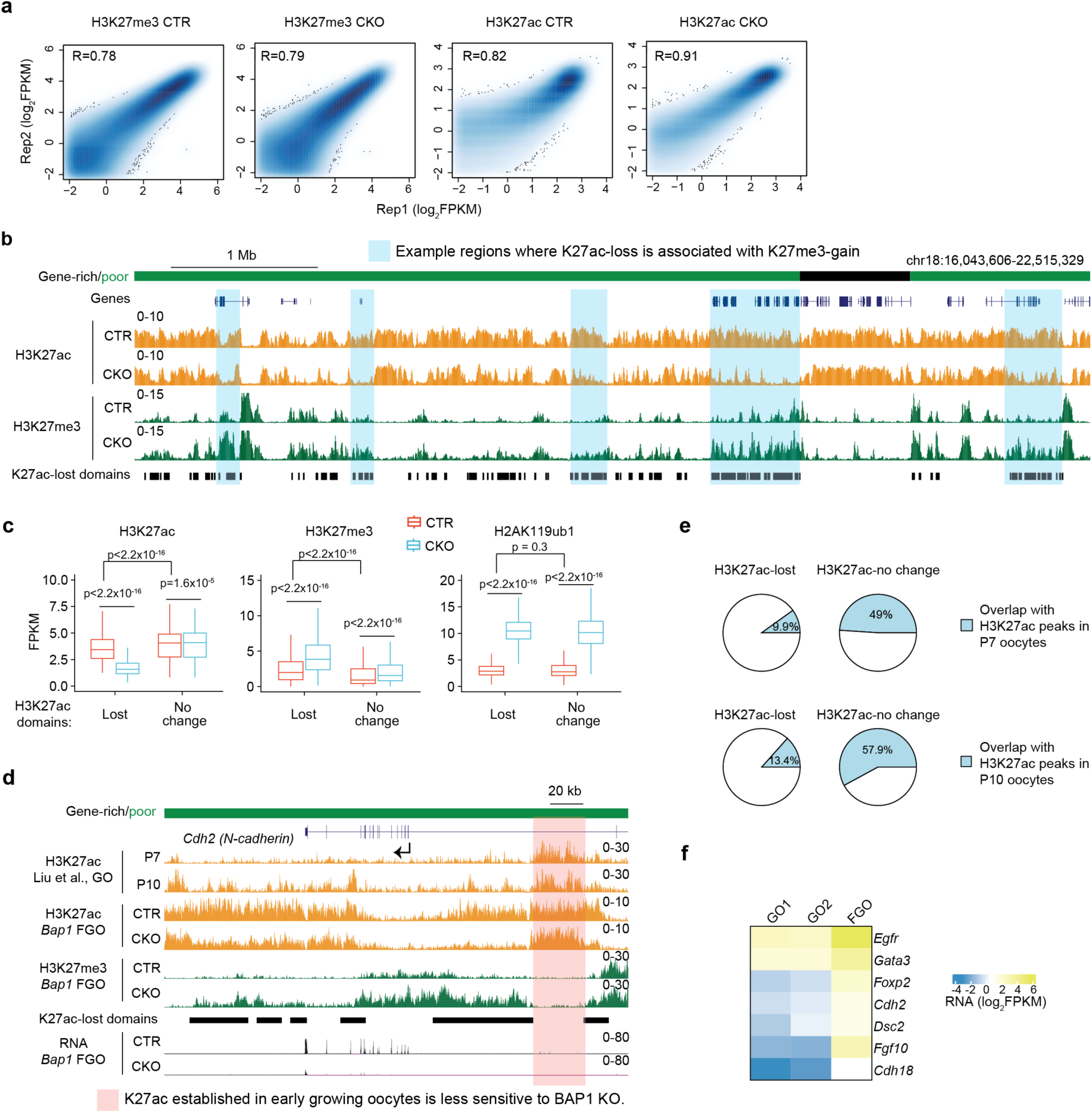
Effects of BAP1 deficiency on H3K27ac and H3K27me3 profiles in FGOs. **a)** Scatter plots showing the reproducibility of H3K27me3 and H3K27ac CUT&RUN signals across biological replicates in FGOs. Pearson correlation coefficients are indicated. **b)** Genome browser view depicting H3K27ac and H3K27me3 signals at the indicated genomic locus. Regions with significantly reduced H3K27ac levels in CKO FGOs (“H3K27ac-lost domains”) are indicated. **c)** Boxplots showing H3K27ac, H3K27me3, and H2AK119ub1 enrichment at “H3K27ac-lost” versus “H3K27ac-no change” domains in FGOs. Boxplot elements: center line = median; box bounds = 25th and 75th percentiles; whiskers = ±1.5× interquartile range. *p* value: two-sided Wilcoxon rank-sum test. **d)** Genome browser view of H3K27ac and H3K27me3 profiles at the *Cdh2* locus. **e)** Proportion of “H3K27ac-lost” and “H3K27ac-no change” domains that overlap with H3K27ac peaks in growing oocytes at postnatal day 7 (P7) and P10. P7/P10 H3K27ac ChIP-seq are from public datasets (Liu *et al*., 2024). **f)** Heatmap depicting expression levels of genes downregulated in *Bap1* CKO FGOs across different stages of wild-type oocyte development: GO1 (early secondary follicle) and GO2 (secondary follicle). RNA-seq data for GO1/GO2/FGO samples are from public datasets (Zhang *et al*., 2020).

**Figure S4.**
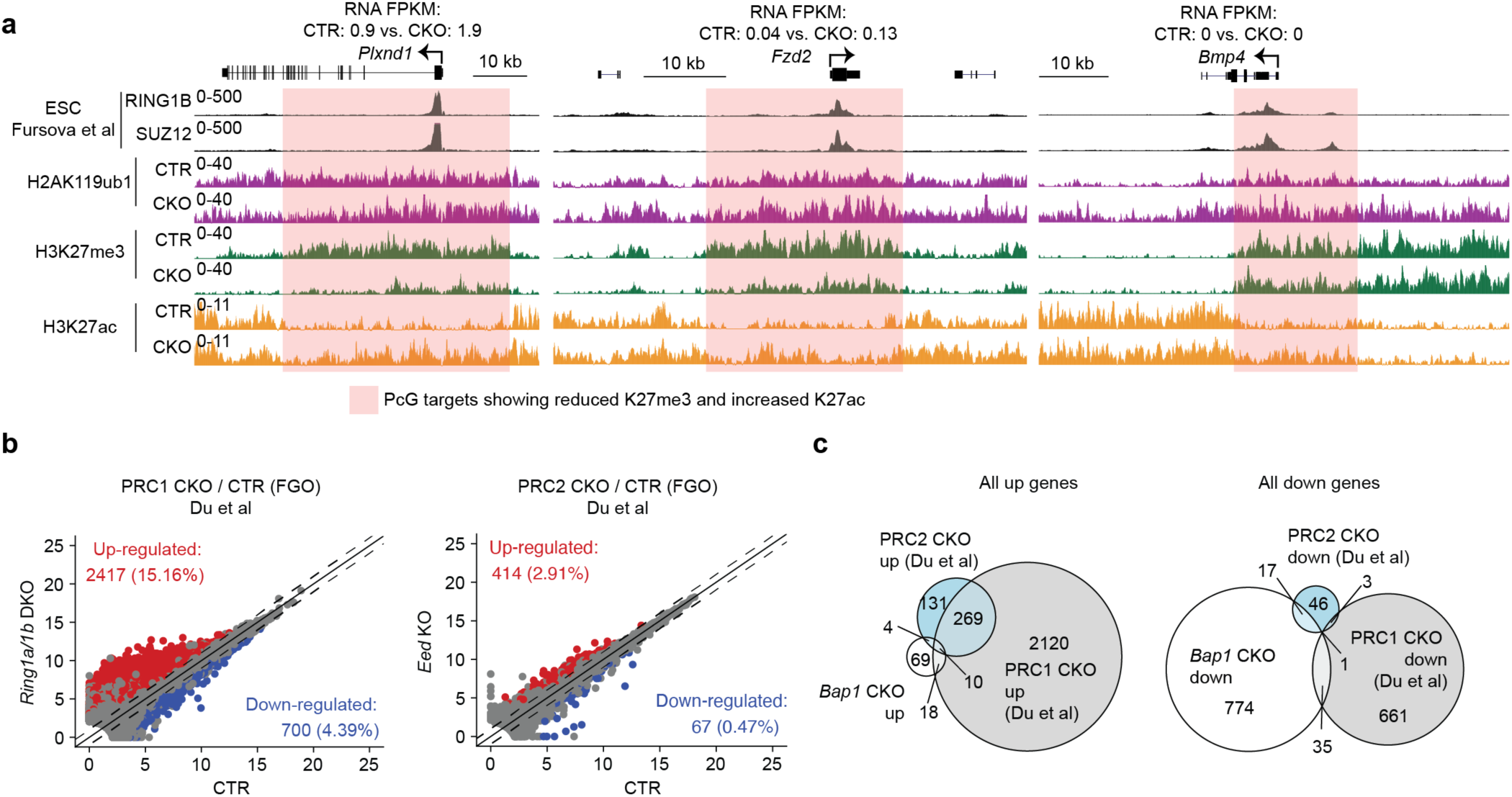
Comparison of PR-DUB, PRC1, or PRC2 knockout effects on the oocyte transcriptome. a) Genome browser views of H2AK119ub1, H3K27me3, and H3K27ac signals at the Polycomb Group (PcG) target loci *Plxnd1, Fzd2,* and *Bmp4*. Public RING1B and SUZ12 ChIP-seq data in embryonic stem cells (ESCs) are included for comparison (Fursova *et al*., 2019). b) Scatter plots comparing gene expression profiles between control (CTR) and PRC1 or PRC2 CKO FGOs. Red and blue dots indicate significantly upregulated and downregulated genes, respectively. Differentially expressed genes were defined by fold change ≥ 2, adjusted *p* < 0.05, and FPKM ≥ 0.5. RNA-seq data for PRC1/2 samples are from public datasets (Du *et al*., 2020). c) Venn diagrams illustrating the overlap of differentially expressed genes among BAP1, PRC1, and PRC2 CKO FGOs.

**Figure S5.**
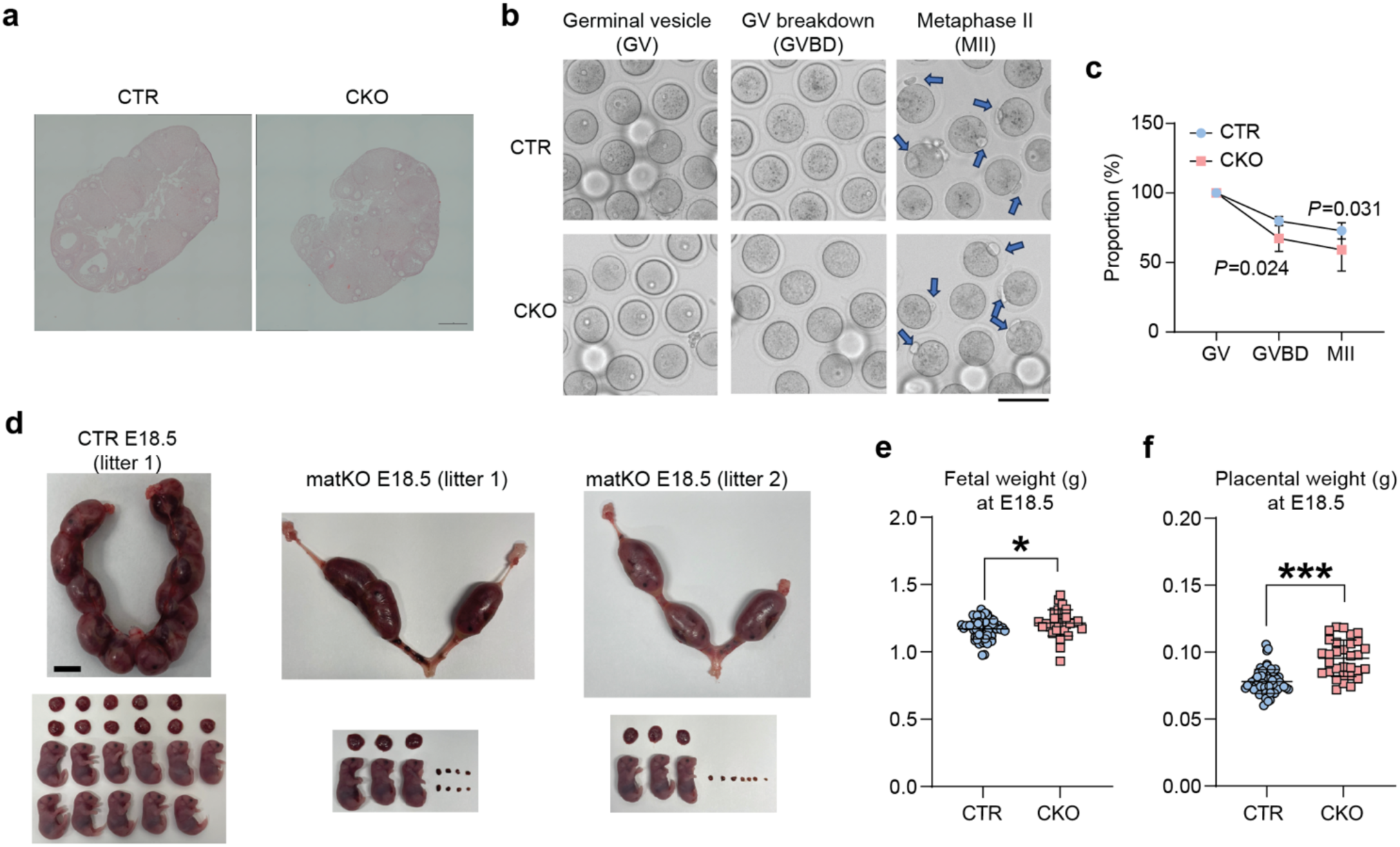
Characterization of BAP1 knockout effects on female fertility. **a)** Histology analyses of ovary sections (n = 2). Scale bar: 0.2 mm. **b)** Brightfield images showing *in vitro* meiotic maturation of *Bap1* CTR and CKO FGOs. Arrows indicate second polar bodies. Scale bar: 100 μm. **c)** Proportions of FGOs reaching the germinal vesicle breakdown (GVBD) and metaphase II (MII) stages. A total of 190 and 165 oocytes were analyzed from CTR (*n* = 4) and CKO (*n* = 4) female mice (9 weeks old), respectively. Data are presented as mean ± SD. *p*-values, chi-square test. **d)** Representative images of uteruses, fetuses, placentae, and resorption sites at E18.5. Scale bar: 1 cm. **e)** Fetal weights at E18.5. CTR: 58 pups from six litters; CKO: 31 pups from eight litters. Data are presented as mean ± SD. \**p* < 0.05, two-sided Student’s *t*-test. **f)** Placental weights at E18.5. Sample sizes match those in panel **g**. Data are presented as mean ± SD. \*\*\**p* < 0.001, two-sided Student’s *t*-test.

**Figure S6.**
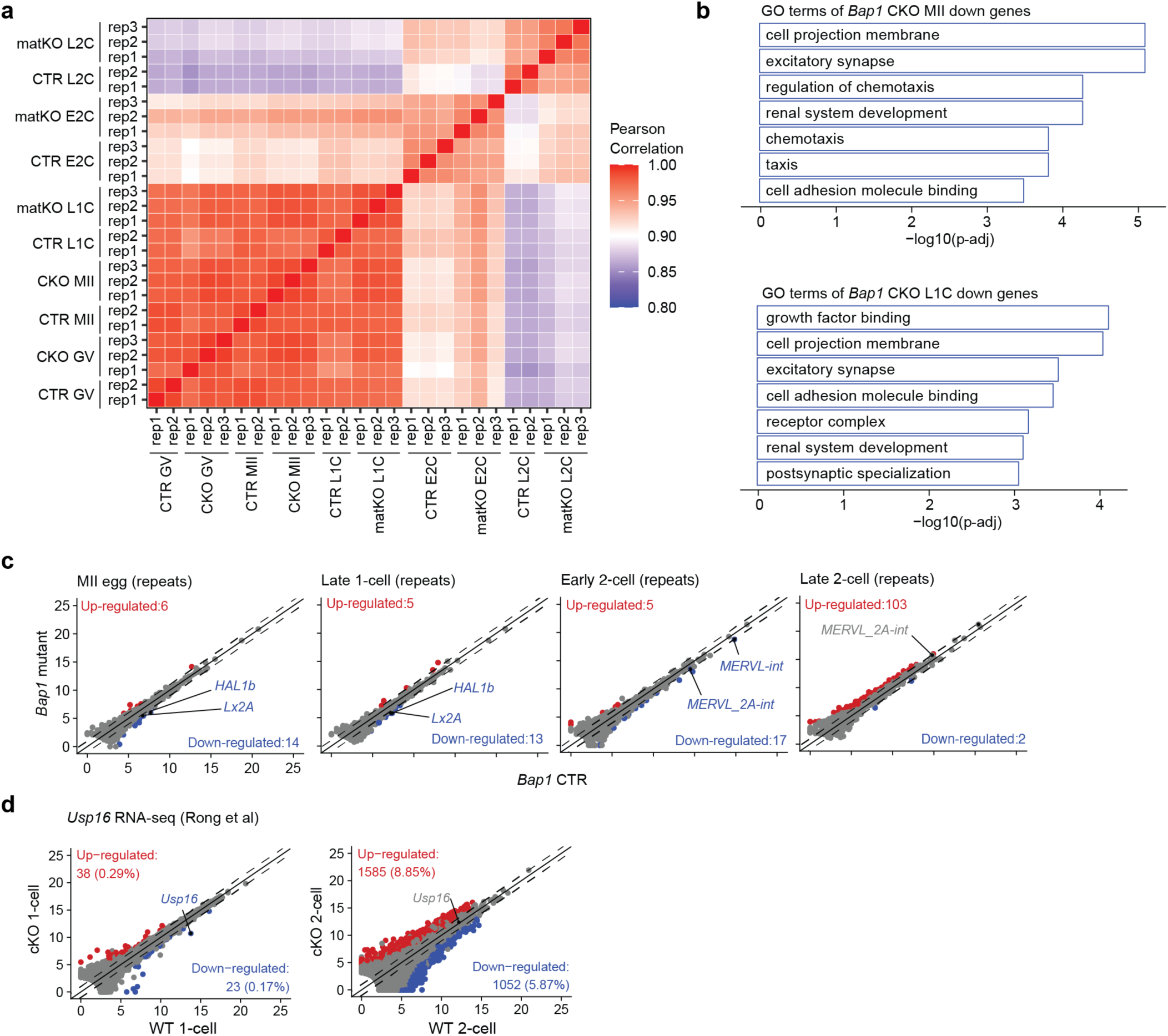
Effects of *Bap1* KO on transcriptomes of oocytes and early embryos. **a)** Heatmap showing reproducibility between biological replicates of total RNA-seq datasets. **b)** Gene ontology (GO) terms enriched among the downregulated genes in *Bap1* CKO MII eggs and late 1-cell (L1C) embryos. **c)** Scatter plot comparing differential expression of repeat element subfamilies in *Bap1* CTR vs. CKO at MII, late 1-cell, early 2-cell, and late 2-cell stages. Red and blue dots indicate significantly upregulated and downregulated repeats in *Bap1* CKO, respectively. Differential expression defined by fold change ≥ 2 and adjusted *p* < 0.05. **d)** Scatter plots comparing gene expression profiles between wild type (WT) and *Usp16* CKO 1-cell and 2-cell embryos. Red and blue dots indicate significantly upregulated and downregulated genes, respectively. Differentially expressed genes were defined by fold change ≥ 2, adjusted *p* < 0.05, and FPKM ≥ 0.5. RNA-seq data are from public datasets (Rong *et al*., 2022).

**Figure S7.**
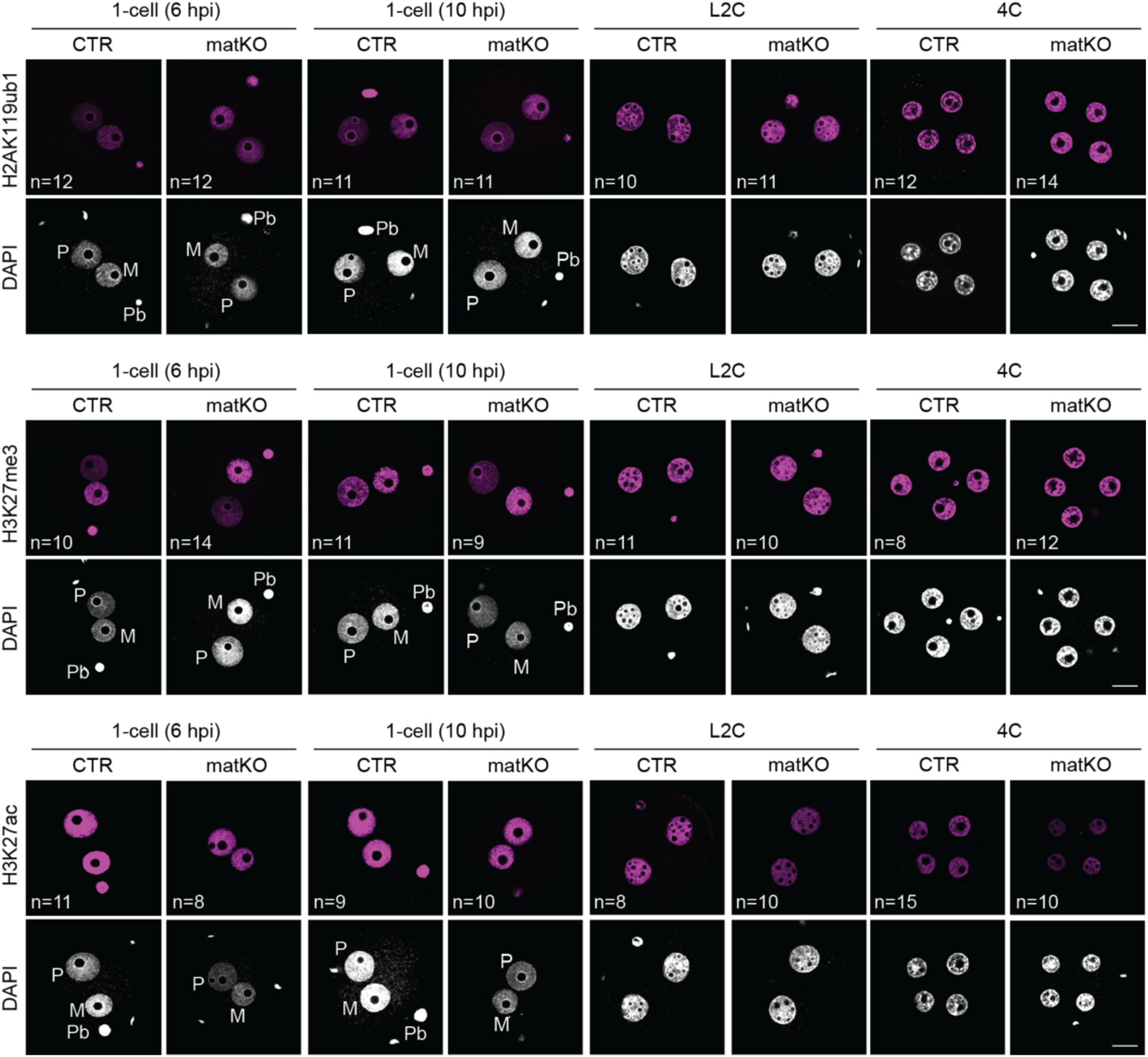
Immunofluorescence images of H2AK119ub1, H3K27me3, and H3K27ac in *Bap1* control (CTR) and maternal KO (matKO) embryos. Number of embryos analyzed are indicated. M: maternal pronuclei; P: paternal pronuclei; Pb: polar body; Scale bar: 20 μm.

**Figure S8.**
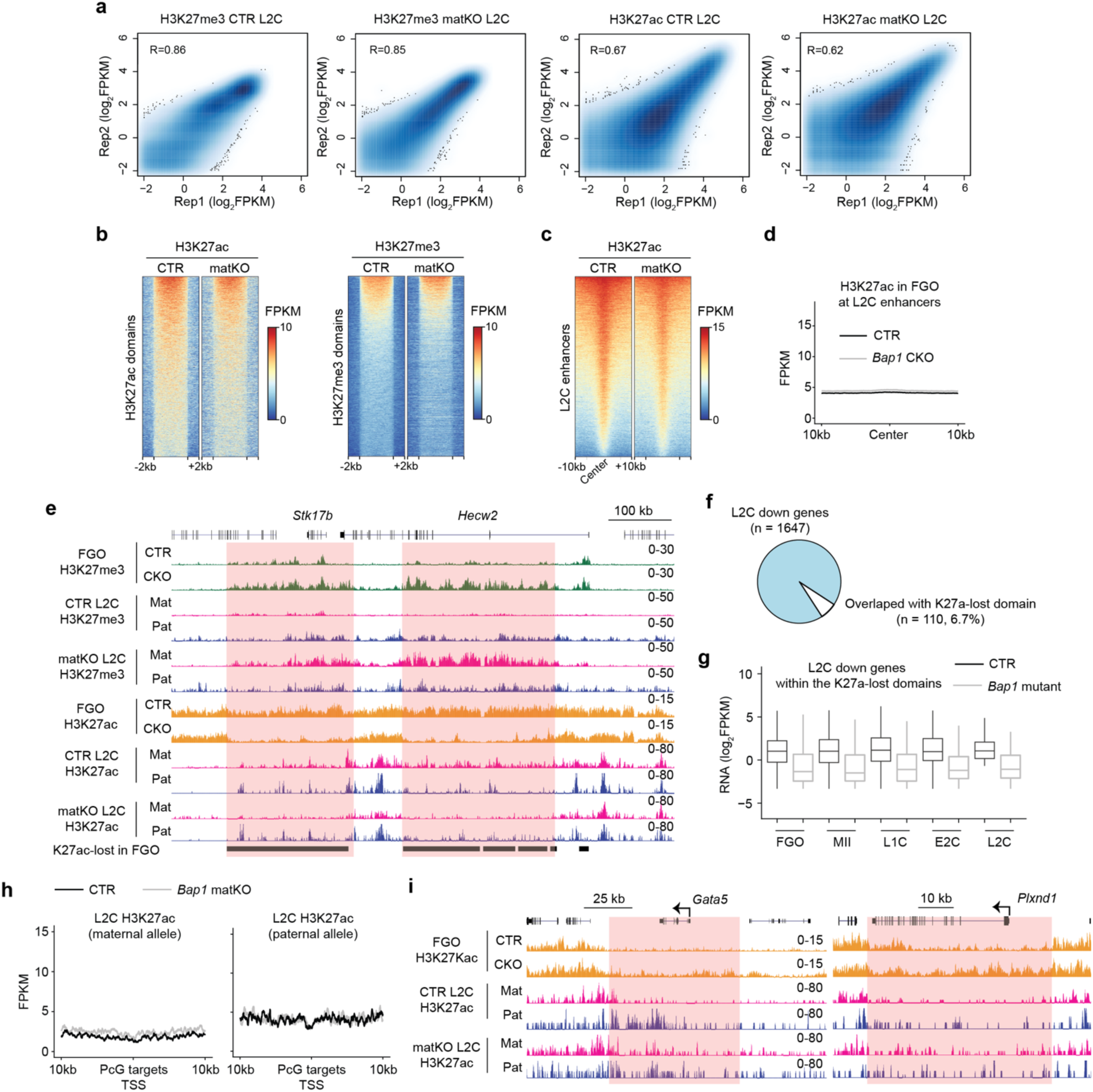
Effects of *Bap1* KO on H3K27me3 and H3K27ac in early embryos. a) Scatter plots showing reproducibility of H3K27me3 and H3K27ac CUT&RUN profiles between biological replicates in late 2-cell (L2C) embryos. Pearson correlation coefficients are indicated. b) Heatmaps showing signal intensity of H3K27ac and H3K27me3 across defined H3K27ac (n = 10,541) and H3K27me3 (n = 9,094) domains in *Bap1* CTR and matKO L2C embryos (see Methods). c) Heatmap depicting signal intensity of H3K27ac at putative enhancers (n=27,899) in *Bap1* CTR and matKO L2C embryos. L2C enhancers were previously defined by distal H3K27ac (Liu *et al*., 2024). d) Metaplot showing H3K27ac signals at L2C enhancers in FGOs. e) Genome browser views of H3K27me3 and H3K27ac profiles in FGOs and L2C embryos at the indicated genomic locus. f) Proportions of downregulated genes in *Bap1* matKO L2C embryos that overlap with the H3K27ac-lost domains. g) Boxplot showing the expression levels of the 110 genes in (**f**) across different developmental stages. Boxplot features: center line indicates the median; box bounds represent the 25th and 75th percentiles; whiskers extend to ±1.5× the interquartile range. h) Metaplot showing allelic H3K27ac levels at a subset of Polycomb Group (PcG) targets in 2-cell embryos. i) Genome browser views of H3K27ac profiles in FGOs and L2C embryos at the indicated PcG targets loci.

